# Tunable molecular tension sensors reveal extension-based control of vinculin loading

**DOI:** 10.1101/264549

**Authors:** Andrew S. LaCroix, Andrew D. Lynch, Matthew E. Berginski, Brenton D. Hoffman

**Affiliations:** Biomedical Engineering, Duke University, Durham NC, USA

## Abstract

Molecular tension sensors have contributed to a growing understanding of mechanobiology. However, the limited dynamic range and inability to specify the mechanical sensitivity of these sensors has hindered their widespread use in diverse contexts. Here, we systematically examine the components of tension sensors that can be altered to improve their functionality. Guided by the development of a first principles model describing the mechanical behavior of these sensors, we create a collection of sensors that exhibit predictable sensitivities and significantly improved performance *in cellulo.* Utilized in the context of vinculin mechanobiology, a trio of these new biosensors with distinct force- and extension- sensitivities reveal that an extension-based control paradigm regulates vinculin loading. To enable the rational design of molecular tension sensors appropriate for diverse applications, we predict the mechanical behavior, in terms of force and extension, of additional 1020 distinct designs.

## Introduction

The ability of cells to generate and respond to mechanical loads is increasingly recognized as a critical driver in many fundamentally important biological processes, including migration (*Doyle et al., 2009; Lo et al., 2000; Pelham and Wang, 1997*), proliferation (*Chen et al., 1997; Provenzano and Keely, 2011*), differentiation (*Engler et al., 2006; Heo et al., 2016; McBeath et al., 2004*), and morphogenesis (*Heisenberg and Bellaiche, 2013; Wozniak and Chen, 2009*). While the mechanosensitive signaling pathways enabling these responses are poorly understood, most are thought to have a common basis: the mechanical deformation of load-bearing proteins (*Cost et al., 2015; Hoffman et al., 2011; Ju et al., 2016*). As such, several technologies for measuring the loads borne by specific proteins in living cells have emerged (*Freikamp et al., 2016a; Hoffman, 2014; LaCroix et al., 2015b*). These biosensors, collectively referred to as molecular tension sensors, leverage the distance-dependence of Förster Resonance Energy Transfer (FRET) to measure the extension of and, if properly calibrated, the forces across a specific protein of interest (*Austen et al., 2013; Freikamp et al., 2016b; Hoffman, 2014; LaCroix et al., 2015a*). For example, using this approach, the tension across vinculin was shown to regulate a mechanosensitive switch governing the assembly/disassembly dynamics of focal adhesions (FAs) (*Grashoff et al., 2010*). While this and several other FRET-based molecular tension sensors provide a critical view into mechanosensitive processes (*Cost et al., 2015; Jurchenko and Salaita, 2015*), fundamental questions regarding the nature and the degree of the mechanical loading of proteins remain. A key limitation has been the inability to create tension sensors with diverse mechanical sensitivities suitable for a wide variety of biological applications (*Freikamp et al., 2016b*).

To date, genetically-encoded molecular tension sensor modules (TSMods), which are incorporated into various proteins to form distinct tension sensors (Figure 1A), have been created without *a priori* knowledge of their mechanical sensitivity. TSMod development has largely relied on a biologically- inspired “guess-and-check” design approach using naturally-occurring extensible polypeptides or protein domains as deformable elements in the FRET-based tension sensors. Furthermore, despite the use of these sensors to study intracellular processes, calibration measurements of their mechanical sensitivity are typically performed *in vitro* using highly precise single molecule techniques. Reported force sensitivities of several *in vitro* calibrated TSMods are 1-6 pN (*Grashoff et al., 2010*), 2-11 pN (*Brenner et al., 2016*), 3-5 pN (*Ringer et al., 2017*), 6-8 pN (*Austen et al., 2015*), or 9-11 pN (*Austen et al., 2015*). However, it is unclear if these ranges are sufficient for diverse mechanobiological studies, and the applicability of these *in vitro* calibrations to sensors that are utilized *in cellulo* has not been verified.

We sought to overcome these limitations by creating new TSMods that do not rely on naturally occurring extensible domains or *in vitro* calibration schemes. These new TSMods consist of an improved Clover-mRuby2 FRET pair connected by unstructured polypeptide extensible domains of various lengths. As the entropy-driven mechanical resistance of unstructured polypeptides can be accurately predicted by established models of polymer extension (*Becker et al., 2010*), the force- and extension-sensitivities can be determined independently of *in vitro* calibration experiments. Using these advancements, we generate a variety of new tension sensors for the FA protein vinculin. These include a version optimized for sensitivity, which shows a 300% increase in performance, as well as a suite of sensors with distinct mechanical sensitivities capable of determining if vinculin loading is subject to extension-based or force-based control. Lastly, we computationally predict the mechanical behavior expected for a variety of unstructured polypeptide-based tension sensors for several common FRET pairs. This resource should allow for the expedited creation and rational design of molecular tension sensors suited for diverse contexts, alleviating a significant limitation in the study of mechanobiology.

## Results

### Creation of TSMods based on synthetic unstructured polypeptides

TSMods for intracellular use consist of two fluorescent proteins (FPs) connected by an extensible domain (Figure 1A). To enable the creation of tension sensors with diverse mechanical sensitivities, we constructed a variety of TSMods using FPs with distinct photophysical properties connected by unstructured polypeptides of various lengths and mechanical properties, as each of these characteristics critically determine the behavior of these sensors (Figure 1B-D). We based our designs on the first calibrated TSMod (*Grashoff et al., 2010*), which is comprised of the mTFP1-Venus FRET pair connected by a flagelliform silk-based polypeptide with the repeated sequence (GPGGA)_8_ and has been used in a variety of tension sensors (*Cost et al., 2015; Jurchenko and Salaita, 2015*).

First, we evaluated the role of the FPs in TSMod function. Reasoning that increases in the unloaded FRET efficiency could potentially increase the dynamic range of the sensor as well as alleviate technical issues with measuring small FRET signals, we sought to increase the FRET efficiency in this state (Figure 1B). To do so, we replaced mTFP1-Venus with the green-red FRET pair Clover-mRuby2 (*Lam et al., 2012*), which exhibits stronger FRET at a given separation distance (Förster radius (*R*_0_) of 5.7 and 6.3 nm, respectively). This simple substitution yielded a 12% higher baseline (unloaded) FRET efficiency that was observed in fixed (Figure 1E, F) and live cells (Figure 1–Figure supplement 1), as well as cell lysates (Figure 1–Figure supplement 2). In addition to their photophysical properties, we considered the effect of the physical structure of the FPs on their performance in TSMods. Although commonly identified by a characteristic beta-barrel structure, FPs also contain short unstructured regions at their termini that likely contribute to the effective mechanical properties of the extensible domains used in TSMods (*Ohashi et al., 2007*). Previous work, and our data (Figure 1–Figure supplement 3), have shown that “minimal” Clover (residues 1 – 227) and mRuby2 (residues 3 – 236) exhibit absorbance and emission spectra indistinguishable from their full-length counterparts (*Austen et al., 2015; Li et al., 1997; Ohashi et al., 2007; Ouyang et al., 2008; Shimozono et al., 2006*). Therefore, to mitigate concerns about FPs affecting the mechanical properties of the extensible domains and further increase the unloaded FRET efficiency, minimal versions of Clover and mRuby2 were used in the construction of all TSMods.

Recent evidence suggests that both the mechanical properties (*Austen et al., 2013; Ringer et al., 2017*) and the length (*Brenner et al., 2016*) of the extensible domain provide viable means by which to tune the mechanical sensitivity of TSMods (Figure 1C, D). Towards this end, we created a variety of TSMods containing extensible domains comprised of either the flagelliform-based (GPGGA)_n_, which is thought to be relatively stiff (*Becker et al., 2003*), or the synthetic (GGSGGS)_n_ which has been characterized as an unstructured polypeptide and has previously been employed as a tunable linker in biochemical sensors (*Evers et al., 2006a*). Analysis of TSMods in cell lysates showed that those with (GGSGGS)_n_ extensible domains exhibit higher FRET efficiencies than those with (GPGGA)_n_ extensible domains of the same length (Figure 1G), suggesting that (GPGGA)_n_-based polypeptides are indeed stiffer, and thus force the FPs apart more readily, than (GGSGGS)_n_-based polypeptides. However, when (GPGGA)_n_ and (GGSGGS)_n_ TSMods were evaluated *in cellulo,* the FRET efficiency versus length relationships were indistinguishable, suggesting the polypeptides are exhibiting identical mechanical properties (Figure 1H). Together, these data demonstrate that factors dictating sensor functionality in the absence of applied load can be environmentally sensitive, and that the behavior of TSMods observed *in vitro* may not be reflective of their behavior *in cellulo.* As such, these results raise concerns about the applicability of calibrations of FRET-based tension sensors performed *in vitro* to sensors that are used in intracellular environments.

**Figure 1.**
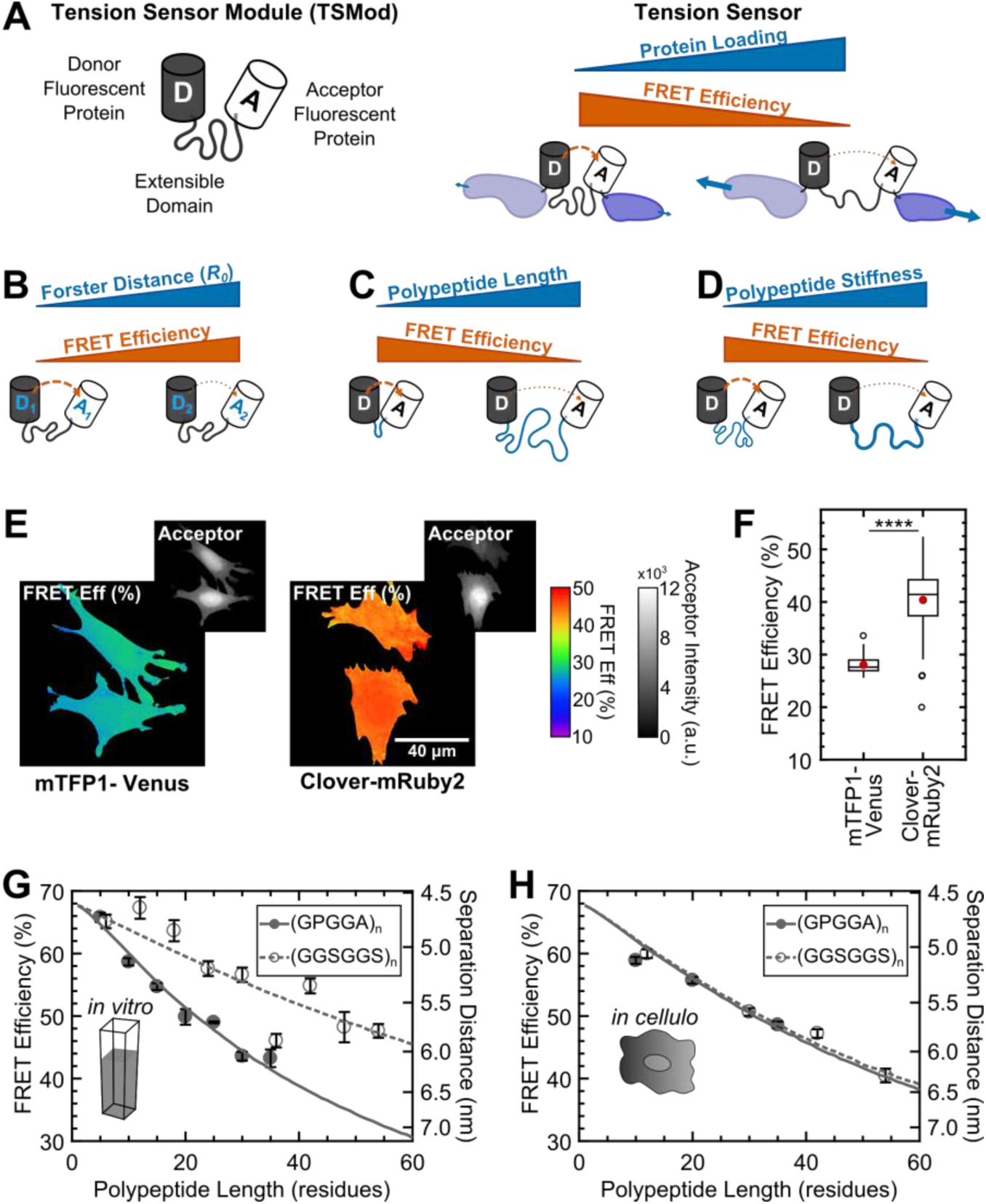
Design and characterization of tunable FRET-based molecular tension sensors. (**A**) Schematic depiction of a generic TSMod and inverse relationship between FRET and force for molecular tension sensors under tensile loading. (**B-D**) TSMod function depends on the Förster radius of the chosen FRET pair (**B**) as well as the length (**C**) and stiffness (**D**) of the extensible polypeptide domain. (**E**) Representative images of soluble mTFP1-Venus and Clover-mRuby2 TSMods expressed in Vin-/- MEFs. (**F**) Quantification of unloaded FRET efficiency for mTFP1-Venus and Clover-mRuby2 TSMods; (n = 53 and 92 cells, respectively); red filled circle denotes sample mean; **** p < 0.0001, Student’s t-test, two-tailed, assuming unequal variances. (**G**) Quantification of FRET-polypeptide length relationship for minimal Clover-mRuby2 TSMods *in vitro;* each point represents data from at least 5 independent experiments; lines represent model fits where *L**P*** is the only unconstrained parameter. (**H**) Quantification of FRET-polypeptide length relationship for minimal Clover-mRuby2 based TSMods *in cellulo;* each point represents at least n = 48 cells from three independent experiments; lines represent model fits where persistence length (*L_P_)* is the only unconstrained parameter. All error bars, s.e.m. **Figure supplement 1** | FRET efficiency measurements are insensitive to fixation and sensor intensity. **Figure supplement 2** | Increase in unloaded FRET efficiency with Clover-mRuby2 sensors *in vitro.* **Figure supplement 3** | “Minimal” FPs exhibit spectral properties indistinguishable from full-length parent FPs.

### A quantitative model describing the mechanical sensitivity of TSMods

As an alternative to TSMod calibration through *in vitro* approaches, we pursued a modeling-based approach for describing the mechanical sensitivities of TSMods. Given that FPs linked by (GGSGGS)_n_ polypeptides are well-described by established models of polymer physics in unloaded conditions (*Evers et al., 2006a*), we developed an analogous model to predict TSMod behavior under load. Briefly, the proposed calibration model incorporates three main aspects of TSMods: (1) the photophysical properties of the FRET pair (Förster radius, *R*_0_), (2) the radii of the FPs (*R_FP_*), and (3) the mechanical response of the extensible domain, which is well-described as a semi-flexible polymer by a persistence length (*L_P_*) and a contour length (*L_C_*) in the framework of the worm-like chain model (*Becker et al., 2010*). This modeling-based approach enables the prediction of the *in cellulo* mechanical response of FRET-based tension sensors by leveraging separate measurements of the *in cellulo L_P_* of the unstructured polypeptide used as the extensible domain. A detailed description of the development and implementation of the model, as well as comparison to other estimates of TSMod behavior are presented in **Figure 2–Supplementary note 1**, which refers to data presented in Figure 2–Figure supplements 1, 2 and **Supplementary tables 1, 2**.

To validate this model, we first investigated its ability to describe the behavior of several types of TSMods in terms of the relationship between FRET and the length of the extensible domain in unloaded conditions. These measurements are critical in that they are used to estimate the mechanics of the extensible domain in terms of its persistence length *L_P_*. To do so, estimates of *R*_0_ and *R_FP_* were obtained from the literature, *L_C_* was directly calculated from the number of acids comprising the extensible domain, and *L_P_* was used as the single adjustable parameter. With only *L_P_* left unconstrained, the model accurately describes the behavior of TSMods containing (GPGGA)_n_ and (GGSGGS)_n_ extensible domains in unloaded conditions in *in vitro* (Figure 1G) and *in cellulo* (Figure 1H) environments with physically reasonable estimates of *L_P_*. Model fits and 95% confidence intervals confirm that *L_P_* estimates for (GPGGA)_n_ and (GGSGGS)_n_ polypeptides are significantly different *in vitro* (0.74 + 0.05 and 0.33 + 0.05 nm, respectively), and collapse to one intermediate value *in cellulo* (0.50 + 0.02 and 0.48 + 0.05 nm, respectively) (see Figure 2-Supplementary table 1 for comparisons). Also, to demonstrate that the literature estimates of *R*_0_ and *R_FP_* were appropriate, we performed a sensitivity analysis, leaving either *R_FP_* or *R*_0_ unconstrained. We observe negligible improvement in fit quality and achieve similar estimates of *L_P_* (Figure 2–Figure supplements 3, 4), validating our approach. Overall, these results demonstrate the functionality of the model to measure the *L_P_* of TSMod extensible domains in unloaded conditions and also suggest that the observed mechanics of the extensible domain can change in different environments, although less-so for (GGSGGS)_n_ polypeptides.

Next, we sought to examine the generalizability of the model as well as validate the ability of the model to describe the behavior of TSMods subject to tensile loads. Therefore, we examined model fits to published fluorescence-force spectroscopy measurements of Cy3-Cy5 dyes linked by (GPGGA)_n_ extensible domains (*Brenner et al., 2016*). Again, with only *L_P_* unconstrained, the model accurately describes the behavior of these TSMod-like constructs in both unloaded conditions (Figure 2A) and under tensile loads (Figure 2B). Importantly, each of these datasets is well-described by the same persistence length (*L_P_* = 1.05 nm) indicating that the same mechanical model is appropriate for describing the behavior of unstructured polypeptides in both unloaded and loaded conditions when both measurements are determined in the same environment. For comparison, we show fits for a range of *L_P_* values from 1.0 to 1.15 nm (lines in Figure 2A, shaded region in Fig 2B). The robustness of these fits to various parameter constraints was also verified (Figure 2–Figure supplement 5). It is important to note that the different estimate of *L_P_* for (GPGGA)_n_ determined from these measurements in Figure 1 is consistent with the idea that *in vitro* calibration should be applied to sensors used in living cells with caution.

**Figure 2.**
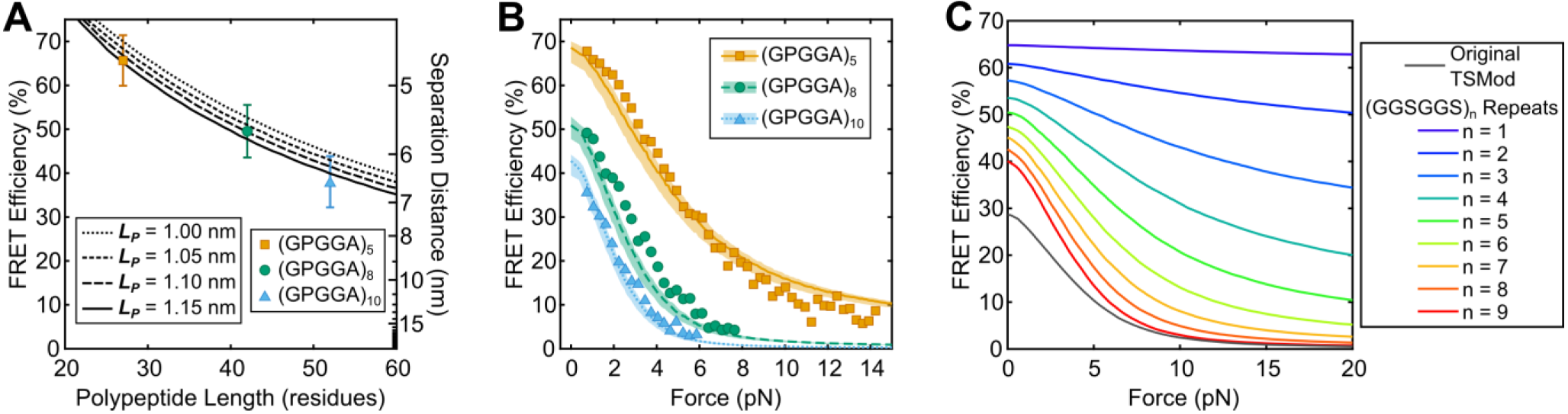
Predicting TSMod calibrations using a biophysical model. (**A, B**) Model descriptions, at various persistence lengths, of FRET- polypeptide length relationship (**A**) and FRET-force responses (**B**) of Cy3 and Cy5 dyes linked by SMCC linker + cysteine modified (GPGGA)_n_ polypeptides; data was digitized based on histograms from (*Brenner et al., 2016*); model parameters *R_0_ = 5.4* nm, *R_FP_* = 0.95 nm, *L_P_* = 1.05 nm (range 1.00 to 1.15 nm); error bars, s.d. (**C**) Model predictions of force sensitivity of TSMods based on Clover-mRuby2 FRET pair and (GGSGGS)_n_ extensible domains in comparison to the original TSMod (*Grashoff et al., 2010*); model parameters *R_0_ = 6.3* nm, *R_FP_ = 2.3* nm, *L_P_* = 0.48 nm. **Supplementary note 1** | Development, validation, and implementation of a computational model describing the mechanical sensitivity of TSMods **Figure supplement 1** | Schematic depiction of biophysical model describing the mechanical sensitivity of TSMods. **Figure supplement 2** | Verification of the proper implementation of a biophysical model describing the mechanical sensitivity of FRET-based TSMods. **Figure supplement 3** | Parameter constraint has minimal effects on measurement of polypeptide persistence length (*L_P_) in vitro.* **Figure supplement 4** | Parameter constraint has minimal effects on measurement of polypeptide persistence length (*L_P_) in cellulo.* **Figure supplement 5** | Parameter constraint has minimal effects on measurement of polypeptide persistence length (*L_P_)* for TSMod-like constructs in unloaded or loaded conditions. **Figure supplement 6** | Experimental and theoretical examinations of other models (*Brenner et al., 2016*) of (GPGGA)_n_ mechanical sensitivity. **Supplementary table 1** | Fit parameters compared to literature estimates of Förster radius (*R*_0_), fluorescent moiety radius (*R_FP_*), and persistence length (*L_P_*). **Supplementary table 2** | Persistence length (*L_P_)* for a variety of published polypeptides.

### A novel approach for predicting the *in cellulo* calibration of TSMods

Together these results suggest a simple model-based calibration scheme in which measurements of extensible domain mechanics (*L*_P_) in unloaded conditions can be utilized to predict TSMod behavior under tensile loading. While our modeling efforts indicate that both (GPGGA)_n_ and (GGSGGS)_n_ polypeptide mechanics are consistent with unstructured polypeptides (Figure 2–Figure supplement 6), we only generate calibration predictions for TSMods containing (GGSGGS)_n_ extensible domains because they are also less sensitive to environmental changes. In the context of the model, the *in cellulo* persistence length of the (GGSGGS)_n_ extensible domain (*L_P_* = 0.48 nm, Figure 1H) is combined with literature estimates of the radii (*Hink et al., 2000*) and photophysical properties (*Lam et al., 2012*) of Clover and mRuby2 to predict the response of (GGSGGS)_n_-based TSMods under applied loads (Figure 2C). This model-based calibration scheme uniquely overcomes the environmental sensitivity of the extensible domain (compare Figure 1G and 1H) by allowing for *in cellulo* measurements of *L_P_* to be used to estimate the mechanical sensitivity of TSMods.

### Optimized tension sensor reveals gradients of vinculin tension across FAs

To determine which extensible domain length will be optimal for measuring tension across vinculin, we evaluated TSMod mechanical sensitivity across different force regimes by calculating the derivative along the FRET-force curve (Figure 2C, Figure 3–Figure supplement 1). Given the original vinculin tension sensor (VinTS) reported average loads of ∼2.5 pN across vinculin that varied from 1 to 6 pN (*Grashoff et al., 2010*), we choose to further investigate the performance of the TSMod containing the nine-repeat extensible domain, as it exhibits the highest sensitivity in this force regime and is capable of capturing the distribution of the loads on vinculin (Figure 3–Figure supplement 1A). This nine-repeat linker also provides a good balance between FRET dynamic range and peak sensitivity (Figure 3–Figure supplement 1B, C). An optimized VinTS (opt-VinTS) was created by genetically inserting this TSMod into vinculin at same site, after amino acid 883, as in the original VinTS design (*Grashoff et al., 2010*).
We assessed the performance of opt-VinTS by evaluating its ability to detect changes in vinculin loading across both subcellular and FA length scales. Vin-/- MEFs expressing either VinTS or opt- VinTS showed indistinguishable cell and FA morphologies (Figure 3A, A’, C, C’, Figure 3–Figure supplement 2). Furthermore, line scans of acceptor intensity across single FAs indicated similar localization of each sensor (Figure 3A’’, A’’’, C’’, C’’’). These findings indicate that the two sensors exhibit identical biologically functionality. At a subcellular length scale, consistent with our previous findings (*Rothenberg et al., 2015*), both VinTS and opt-VinTS report highest loads (lowest FRET efficiency) in the cell periphery, and no appreciable tensile loading of vinculin in the cell center (Figure 3B, D). Based on previous reports of gradients of vinculin loading within individual FAs (*Sarangi et al., 2017*) and a skewed distribution of mechanical stresses at the cell-substrate interface (*Blakely et al., 2014; Legant et al., 2013; Morimatsu et al., 2015; Plotnikov et al., 2012*), we expected to see similar distally-skewed vinculin tensions. Such gradients are difficult to discern in peripheral FAs of cells expressing the original tension sensor (Figure 3B’, B’’, B’’’). However, individual FAs in the periphery of cells expressing opt-VinTS revealed striking gradients of vinculin tension across single FAs (Figure 3D’, D’’, D’’’). Quantification of FRET efficiency change across length-normalized FAs (Figure 3B’’’, D’’’) reveals an almost 300% improvement in the performance opt-VinTS when compare the original design. In total, these results show that, as predicted by the model, opt-VinTS is significantly more sensitive than the original VinTS.

**Figure 3.**
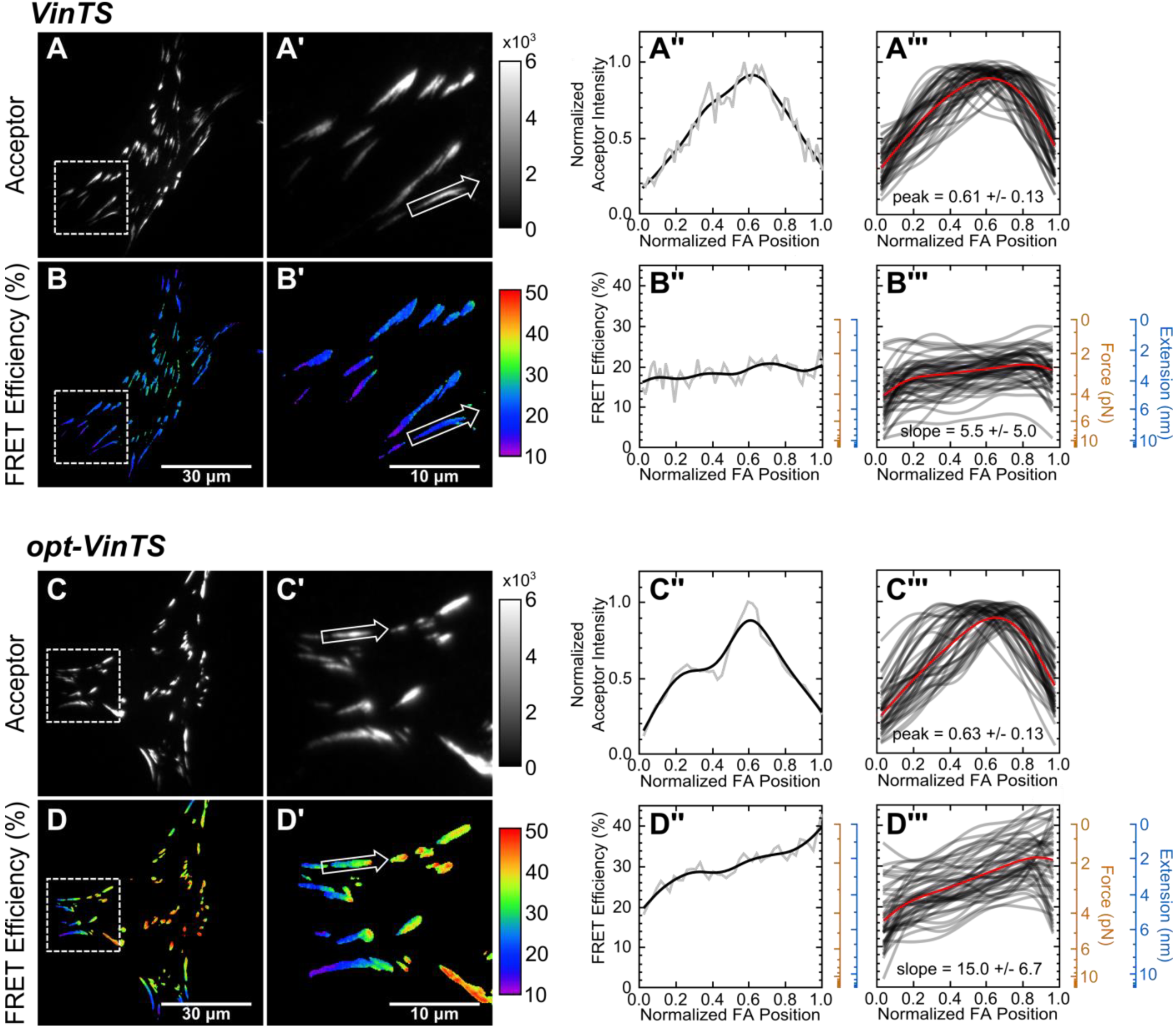
Optimized tension sensor reveals sub-FA gradients in vinculin tension. (**panels A and B**) Representative images of subcellular distribution of VinTS (*Grashoff et al., 2010*) (**A, A’**) along with representative (**A’’**) and aggregate (**A’’’**) line scans of single FAs of size > 0.5 mm^2^ and axis ratio > 1.5; corresponding masked FRET images (**B, B’**) and line scans (**B’’, B’’’**); n = 55 FAs from n = 11 cells from three independent experiments; averaged intensity and FRET profiles in red. (**panels C and D**) As in panels **A** and **B**, but with optimized opt-VinTS construct; n = 49 FAs from n = 10 cells from three independent experiments.
Figure supplement 1 | Selection of optimal (GGSGGS)_n_ extensible domain length in a Clover-mRuby2 based TSMod for measuring ∼1-6 pN loads borne by vinculin.
Figure supplement 2 | FA morphologies, cell morphologies, and sensor localization to FAs are indistinguishable between different versions of VinTS.

### Vinculin loading is subject to an extension-based control mechanism

A central premise of mechanotransduction, the process by which cells sense and respond to mechanical stimuli, is that applied loads induce conformational changes in mechanosensitive proteins, leading to biochemically distinct functions. However, it is unknown whether the forces or the extensions experienced by proteins mediate the activation of mechanosensitive signaling pathways. Experimental evaluation of this molecular-scale question has been challenging because force and extension are inherently linked. For example, in the case of molecular tension sensors, the force-extension relationship for the extensible domain is monotonic, so any given force corresponds to a unique extension (Figure 4–Figure supplement 1). Therefore, to determine whether conserved forces or extensions mediate vinculin loading, we created a trio of vinculin tension sensors with extensible domains comprised of five, seven, or nine repeats of (GGSGGS)_n_. As each sensor has a unique force-extension curve, the application of equivalent forces to the three constructs will result in three distinct extensions, and vice versa (Figure 4–Figure supplement 1). Cells expressing equivalent amounts of each sensor spread and formed FAs normally (Figure 4A-D, Figure 3–Figure supplement 2). Using the *in cellulo* calibration predictions described above (shown in Figure 2C), measured FRET efficiencies (Figure 4E-H) were converted to sensor forces (Figure 4I-L) and extensions (Figure 4M-P). Intriguingly, we observed similar distributions of extension (Figure 4P), and distinct distributions of tensile forces (Figure 4L) in the various sensors. This result strongly suggests that loads across vinculin are governed by an extension-based control rather than the more commonly assumed force-based control paradigm.

**Figure 4.**
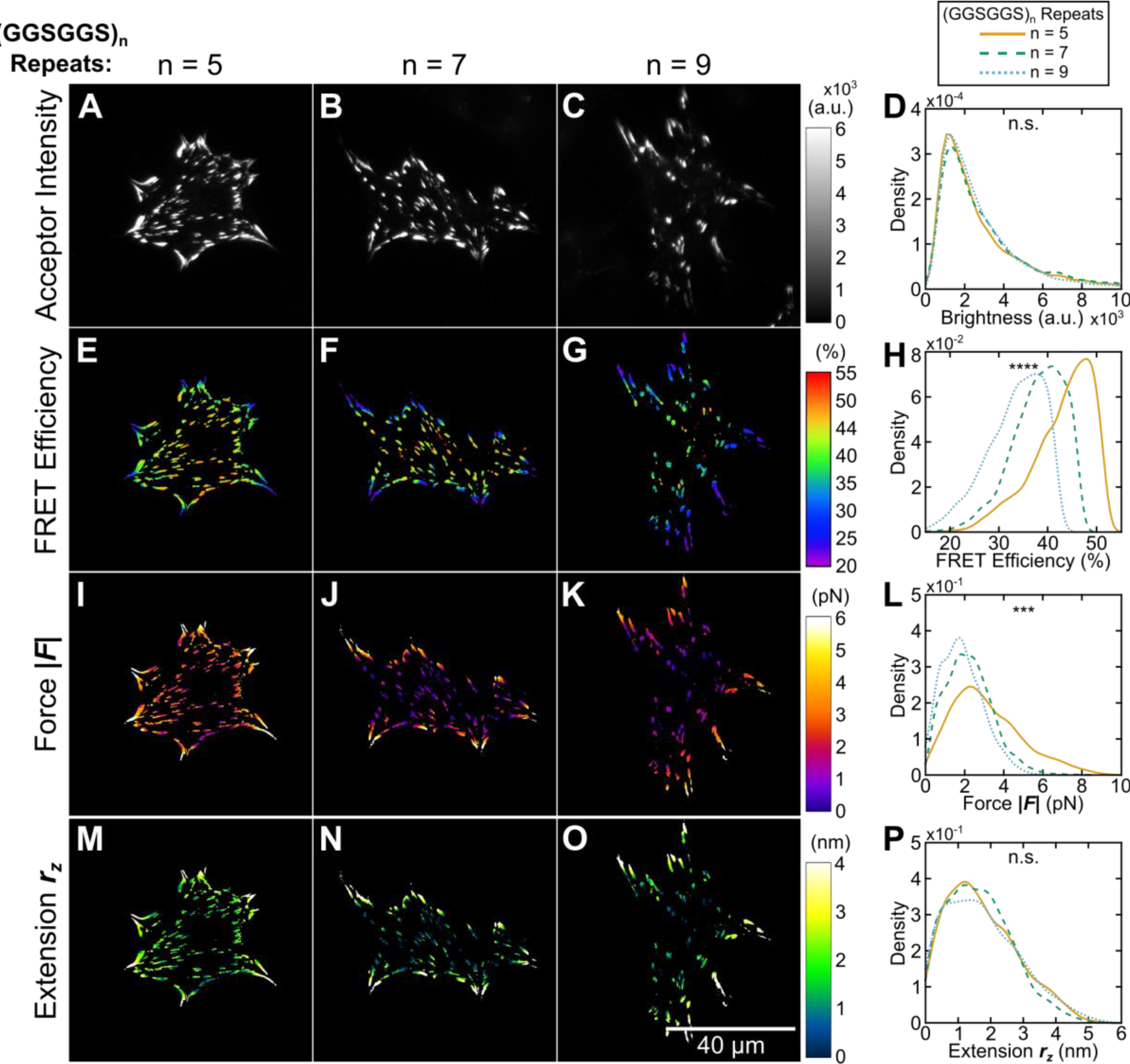
Using tension sensors with distinct mechanical sensitivities to test force-based versus extension-based control of vinculin loading. (**A-C**) Representative images of the localization of a trio of vinculin tension sensors to FAs. (**D**) Normalized histograms of acceptor intensities at FAs is indistinguishable between the three sensors. (**E-G**) Representative images of masked FRET efficiency and (**H**) normalized histograms of average FA FRET reported by each sensor. (**I-K**) Representative images of forces and (**L**) normalized histograms of average vinculin tension in FAs reported by each sensor. (**M-O**) Representative images of extension and (**P**) cumulative distributions of average vinculin extension in FAs reported by each sensor; n = 54, 40, 33 cells and n = 5618, 4837, 4195 FAs for (GGSGGS)5,7,9 extensible domains, respectively; data pooled from three independent experiments; *\*\*\*\** p < 0.0001, *** p < 0.001, n.s. not significant (p ≥ 0.05), ANOVA. See **Supplementary Table 5** for exact p-values and multiple comparisons test details. **Supplementary note 1** | Examining extension-based control of vinculin loading with a structural model of a FA Figure supplement 1 | Schematic depiction of force-extension relationships and potential force- and extension-control paradigms. Figure supplement 2 | Various structural arrangements within FAs could lead to force-based or extension-based control.

To gain insight into the physical origins of force- versus extension-controlled loading of proteins within FAs, we examined how forces and extensions propagate through a simple structural model of a FA (see **Figure 4–Supplementary note 1** for details and more comprehensive discussion of model results). Briefly, the structural model is comprised of various numbers of two distinct elements, which can be thought of as mechanically-dominant proteins or protein complexes within FAs. A sensor element (subscript ⁈S’) and an alternative linker element (subscript “L”) are arranged in two layers (Figure 4–Figure supplement 2A) meant to simulate the stratified organization of FAs (*Kanchanawong et al., 2010*). By comparing the relative variances in forces and extensions observed across sensor elements within various arrangements (Figure 4–Figure supplement 2B), we examined whether a force-controlled or extension-controlled loading of the sensor element would be observed following a bulk force or extension input to the entire structure, and whether this depended on either the relative molecular abundance or the relative stiffness of each element (Figure 4–Figure supplement 2C). Regardless of the relative abundance of the elements or their respective stiffness a force input to the entire structure always resulted in force-based control within the sensor elements. In contrast, an extension input to the entire structure, as might arise due to defined myosin motor step size (*Murphy et al., 2001*) or actin polymerization (*Peskin et al., 1993*), gave rise to both extension-controlled and force-controlled regimes in the sensor elements. The extension-controlled loading of the sensor element is more strongly observed when the sensor element is relatively soft and/or in low abundance, otherwise a force-controlled system is predicted (Figure 4–Figure supplement 2D). Furthermore, in the extension-controlled regime, this simple model also predicts the experimentally-observed linear relationship between sensor element stiffness and the force borne by the three sensor elements (Figure 4–Figure supplement 2E, asterisk). Together, these results demonstrate that protein extension, instead of applied force, might be a key mechanical variable in some mechanosensitive processes.

### Roadmap for future TSMod design

By expanding the simulated parameter space, the calibration model can also be used to predict the *in cellulo* mechanical sensitivity of a wide variety of potential TSMod designs. Specifically, as each model parameter corresponds to an alterable variable in sensor design (*R_0_* = FRET pair, *L_P_* = composition of extensible domain, *N* = length of extensible domain), we can bypass the need to iteratively “guess and check” the performance of new sensors, and, instead, predict the performance of unstructured polypeptide-based tension sensors *in silico*. Since our measurements and modeling efforts indicate that both force and extension might be key mechanical variables in different contexts, we report the predicted mechanical responses for simulated sensors in terms of both force and extension. The predicted relationships between force, extension, and FRET for a single sensor can be concisely described by three metrics as depicted in Figure 5A-C: (1) a FRET dynamic range (*ΔFRET*), which is defined as the change in FRET efficiency from an unloaded state to an experimentally-determined 5% noise floor; (2) a target force (*F_target_*), which indicates the midpoint of force range a sensor is functional, and is defined as *F_target_* = *ΔF/2;* and (3) a target extension (*r_z_,_target_*), which is analogous to target force. Examining the predicted *ΔFRET, F*_target_, and *r_ztarget_* for a variety of Clover-mRuby2 TSMods containing extensible domains of various lengths and compositions, we generate a “roadmap” for future Clover-mRuby2 sensor design (Figure 5D-F, see Figure 2-Supplementary table 2 for a list of reported polypeptide mechanical properties justifying the range over which simulations were performed). Additional roadmaps were generated for other commonly used FRET pairs (Figure 5–Figure supplement 1). With these roadmaps as a guide, the rational design and implementation of future tension sensors with diverse and *a priori* specified properties is now possible.

**Figure 5.**
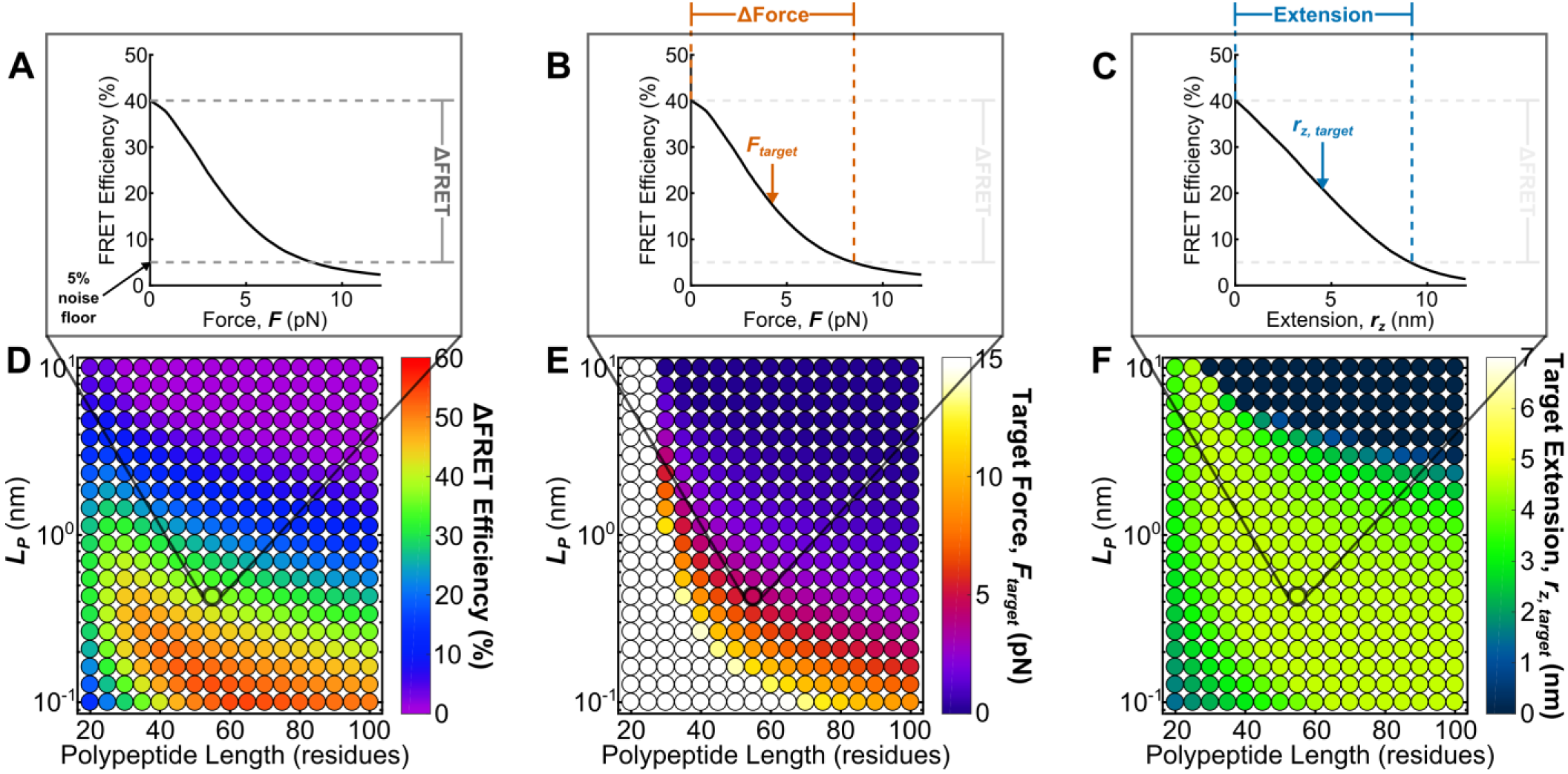
Roadmap to enable the rational design of FRET-based molecular tension sensors. (**A-C**) Representative plots of relationships between FRET efficiencies, forces, and extensions reported by a single sensor, highlighting *ΔFRET* (**A**), *ΔForce* (**B**), and polypeptide extension *r_z_* (**C**) required to stretch a sensor from its resting state to the 5% FRET noise floor. (**D-F**) Parameter space highlighting the predicted *ΔFRET* at the 5% noise floor (**D**) as well as the target force *F_target_* (**E**) and target extension *r_z,target_* (**F**) predicted for a variety of Clover-mRuby2-based sensors with a variety of polypeptide lengths (x-axis), and polypeptide mechanical properties (y-axis). Each point represents a single potential TSMod design. Figure supplement 1 | Roadmaps to guide the rational design of FRET-based molecular tension sensors for some commonly used FRET-pairs.

## Discussion

In this study, we created and characterized a suite of molecular tension sensors with improved, predictable, and tunable force sensitivities. Through these improvements, which included a switch to the Clover-mRuby2 FRET pair, the use of ‘minimal’ FPs, and the modulation of the length and composition of the unstructured polypeptide-based extensible domain, we encountered surprising context-dependent mechanical responses in various TSMods. These context-dependent responses question the applicability of *in vitro* (single molecule-based) calibrations to sensors that are almost exclusively utilized *in cellulo*. To overcome this obstacle, we developed and validated a first-principles model which predicts TSMod mechanical sensitivity using *in cellulo* measurements of the mechanical properties of unstructured polypeptides used as the extensible domain. Then, guided by this model, we created new sensors that provide the next generation of calibrated TSMods for application to diverse mechanobiological investigations.

Initial experiments employing these new sensors highlight their enhanced ability to detect asymmetric distributions of molecular tension within single FAs. Similar tension asymmetries have been observed external to the cell using both high resolution traction force microscopy (*Legant et al., 2013; Plotnikov et al., 2012*) and extracellular tension sensors (*Blakely et al., 2014; Morimatsu et al., 2015*). Intracellularly, gradients in vinculin tension have been reported in cells adhered to micropost arrays, although tension asymmetries were mainly attributed to the presence of the discontinuous substrates (*Sarangi et al., 2017*). We show that these asymmetric molecular loads, high at the distal tips and low at the proximal edge of FAs are transmitted through vinculin even on continuous substrates. Tension asymmetries in continuous systems are likely due to interactions between vinculin and load-transmitting intracellular binding partners, such as F-actin.

This next generation of calibrated TSMods also enabled the investigation of a question that was previously technically inaccessible: are the forces across or extensions of proteins subject to cellular control? The observation that vinculin loading is characterized by repeatable patterns of extension suggests that protein extension, rather than force, may govern the activation of vinculin-dependent and possibly other mechanosensitive signaling pathways. Extrapolated to longer length scales, this result agrees well with reports that cells induce similar strains within extracellular environments of differing stiffness (*Saez et al., 2005*) and that strain determines the activation of touch-sensitive channels in *in vivo* models (*Eastwood et al., 2015*). However, while vinculin appears to be under extension-control in 2D culture systems we note that the simple FA structural model demonstrates that the extent of force-controlled versus extension-controlled protein loading is highly sensitive to the relative abundance and stiffness of various proteins within the bulk structure. Thus, it seems likely that other load-bearing proteins might be subject to either type of control or that a specific protein in distinct cellular contexts could switch control modalities. The prediction of >1000 possible tension sensor designs should allow the creation of tension sensors for any need in either extension- or force-based control paradigms.

In sum, this work provides the biophysical foundation for understanding molecular tension sensor function and delivers a suite of *in cellulo*-calibrated sensors whose distinct and predictable mechanical sensitivities can be leveraged to gain unique molecular understanding of the role of mechanical forces and extensions in biological systems. These advancements should expedite deployment of molecular tension sensors in diverse biological contexts where mechanical cues and cellular force generation have long been thought to play critical, but unexplored, roles.

## Materials and methods

### Cell culture and transfection

Vinculin -/- MEFs (kindly provided by Dr. Ben Fabry and Dr. Wolfgang H. Goldmann (*Mierke et al., 2010*), Friedrich-Alexander-Universitat Erlangen-Nurnberg) were maintained in high-glucose DMEM with sodium pyruvate (D6429, Sigma Aldrich, St. Louis, MO) supplemented with 10% FBS (HyClone, Logan, UT), 1% v/v non-essential amino acids (Invitrogen, Carlsbad, CA), and 1% v/v antibiotic-antimycotic solution (Sigma Aldrich). HEK293 cells were maintained in high-glucose DMEM (D5796, Sigma Aldrich) supplemented with 10% FBS (HyClone) and 1% v/v antibiotic-antimycotic solution (Sigma Aldrich). Cells were grown at 37 °C in a humidified 5% CO2 atmosphere. Cells were transfected at 50-75% confluence in 6-well tissue culture plates using Lipofectamine 2000 (Invitrogen) following the manufacturer’s instructions.

### Generation of TSMods and vinculin tension sensor constructs

Constructs were created from the previously generated pcDNA3.1 mTFP1 (*Ai et al., 2006; Grashoff et al., 2010*) and pcDNA3.1 Venus (A206K) (*Grashoff et al., 2010; Nagai et al., 2002*) as well as pcDNA3.1 Clover (Addgene 40259) and pcDNA3.1 mRuby2 (Addgene 40260) (*Lam et al., 2012*). Minimal versions of single FPs were generated via Polymerase Chain Reaction (PCR) and inserted into pcDNA3.1 via EcoRI/NotI digestion and subsequent ligation (T4 DNA Ligase, NEB, Ipswich, MA). Specifically, creation of minimal FPs involved deletion of the 11 C-terminal residues in mTFP1 and Clover, and the first and second N-terminal residues in Venus and mRuby2 after the start codon. Oligonucleotide primers used to generate full-length and minimal versions of mTFP1, Venus A206K, Clover, and mRuby2 are detailed in **Supplementary file 1**.

The FP component fragments of the mTFP1-Venus and Clover-mRuby2 TSMods were derived from pcDNA3.1 TS module (*Grashoff et al., 2010*) and pcDNA3.1-Clover-mRuby2 (Addgene 49089) or the minimal FP variants described above. The extensible (GPGGA)_n_ and (GGSGGS)_n_ extensible domains were derived from pcDNA3.1 TS module (*Grashoff et al., 2010*) and pET29CLY9 (Addgene 21769) (*Evers et al., 2006a*), respectively. Gibson assembly was used to construct TSMods containing a given FRET pair and extensible domain from three fragments: (1) vector backbone and donor FP (complementary regions: 5’-ampicillin gene, 3’-donor FP), (2) extensible domain region (complementary regions: 5’-donor FP, 3’-acceptor FP), and (3) vector backbone and acceptor FP (complementary regions: 5’-acceptor FP, 3’-ampicillin gene). Primers used to generate the extensible domain region in this reaction scheme were designed to nonspecifically target the repetitive extensible domain sequence, thereby generating extensible domains of various lengths. Following assembly and transformation into DH5α competent cells, single colonies were assayed for extensible domain length by DNA sequencing. Oligonucleotide primers used to generate TSMods are detailed in **Supplementary file 1**.

All variants of the vinculin tension sensor were derived from pcDNA3.1 Vinculin TS (*Grashoff et al., 2010*). In a cloning strategy analogous to that described above for the TSMods, Gibson assembly techniques were used to assemble vinculin tension sensors containing various minimal Clover-mRuby2 TSMods based on three fragments: (1) vector backbone and vinculin head domain residues 1 - 883 (complementary regions: 5’-ampicillin gene, 3’-Clover), (2) TSMod with desired (GGSGGS)_n_ extensible domain (complementary regions: 5’-Clover, 3’-mRuby2), and (3) vector backbone and vinculin tail domain residues 884 - 1066 (complementary regions: 5’-mRuby2, 3’-ampicillin gene). Again, the assembled DNA fragments were transformed into DH5α competent cells and extensible domain length was verified for single colonies by DNA sequencing. Oligonucleotide primers used to generate vinculin tension sensors are detailed in **Supplementary file 1**.

### FRET imaging

For cell imaging, glass coverslips (#1.5, Bioptechs, Butler, PA) placed in reusable dishes (Bioptechs) were coated overnight at 4 °C with 10 μg/mL fibronectin (Fisher Scientific, Pittsburgh, PA). Transfected vinculin -/- MEFs expressing a given tension sensor construct were then trypsinized, transferred to the prepared glass-bottom dishes at a density 50,000 cells per dish, and allowed to spread to 4 h. For fixed experiments, samples were then rinsed quickly with PBS, and fixed for 10 min with 3.7% methanol-free paraformaldehyde (Electron Microscopy Sciences, Hatfield, PA). For live experiments, growth media was exchanged, at least 1 h before imaging, for imaging media - Medium 199 (Life Technologies, 11043) supplemented with 10% FBS (HyClone), 1% v/v non-essential amino acids (Invitrogen), and 1% v/v antibiotic-antimytotic solution (Sigma Aldrich). Live cell imaging was performed for up to 30 min at 37 °C (Stable Z system, Bioptechs).

All imaging was performed on an Olympus IX83 inverted epifluorescent microscope (Olympus, Center Valley, PA) equipped with a LambdaLS 300W ozone-free xenon bulb (Sutter Instruments, Novato, CA), a sCMOS ORCA-Flash4.0 V2 camera (Hamamatsu, Japan), motorized filter wheels (Sutter Instruments 10-3), and automated stage (H117EIX3, Prior Scientific, Rockland, MA). Image acquisition was controlled by MetaMorph Advanced software (Olympus). Samples were imaged at 60X magnification (Olympus, UPlanSApo 60X/NA1.35 objective, 108nm/pix), using a three-image sensitized emission acquisition sequence (*Chen et al., 2006*). The filter set for FRET imaging of mTFP1-Venus sensors includes mTFP1 excitation (ET450/30x, Chroma, Bellows Falls, VT), mTFP1 emission (Chroma, ET485/20m), Venus excitation (Chroma, ET514/10x), and Venus emission (FF01-571/72, Semrock, Rochester, NY) filters, and a dichroic mirror (Chroma T450/514rpc). Images of mTFP1-Venus sensors were acquired in, sequentially, the acceptor channel (Venus excitation, Venus emission, 1000 ms exposure), FRET channel (mTFP1 excitation, Venus emission, 1500 ms exposure), and donor channel (mTFP1 excitation, mTFP1 emission, 1500 ms exposure). For Clover-mRuby2 sensors, we utilized the FITC and TRITC filters from the DA/FI/TR/Cy5-4X4 M-C Brightline Sedat filter set (Semrock), which provided efficient Clover excitation (FF02-485/20), Clover emission (FF01-525/30), mRuby2 excitation (FF01-560/25), and mRuby2 emission (FF01-607/36) filters, and appropriate dichroic mirror (FF410/504/582/669-Di01) for FRET imaging. Images of Clover-mRuby2 sensors were acquired in, sequentially, the acceptor channel (mRuby2 excitation, mRuby2 emission, 1500 ms exposure), FRET channel (Clover excitation, mRuby2 emission, 1500 ms exposure), and donor channel (Clover excitation, Clover emission, 1500 ms exposure).

### Quantitative FRET efficiency measurements from 3-cube FRET imaging

FRET was detected through measurement of sensitized emission (*Chen et al., 2006*) and subsequent calculations were performed on a pixel-by-pixel basis using custom written code in MATLAB (Mathworks, Natick, MA) (*Rothenberg et al., 2015*). Prior to FRET calculations, all images were first corrected for uneven illumination, registered, and background-subtracted. Spectral bleed-through coefficients were determined through FRET-imaging of donor-only and acceptor-only samples (i.e. cells expressing a single donor or acceptor FP). Donor bleed-through coefficients (*dbt*) were calculated for mTFP1 and Clover as:

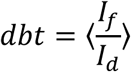

where *I_f_* is the intensity in the FRET-channel, *I_d_* is the intensity in the donor-channel, and data were binned by donor-channel intensity. Similarly, acceptor bleed-through coefficients (*abt*) were calculated for Venus and mRuby2 as:

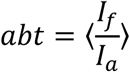

where *I_a_* is the intensity in the acceptor-channel, and data were binned by acceptor-channel intensity. To correct for spectral bleed-through in experimental data, corrected FRET images (*F*_c_) were generated following the equation:

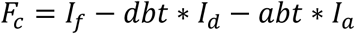

After bleed-through correction, FRET efficiency was calculated following the equation:

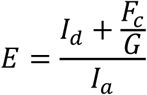

where *G* is a proportionality constant that describes the increase in acceptor intensity (due to sensitized emission) relative to the decrease in donor intensity (due to quenching) (*Chen et al., 2006*). This constant depends on the specific FRET pair used, imaging system, and image acquisition settings, and was calculated for both mTFP1-Venus and Clover-mRuby2 biosensors through imaging donor-acceptor fusion constructs of differing but constant FRET efficiencies. See **Supplementary file 2** for bleed-through and *G* coefficients.

Wherever possible, image analysis was standardized using custom-written Matlab software. Analysis parameters (**Supplementary file 2**) and thresholds for image segmentation were maintained across multiple days of experiments of the same type. For all TSMod and VinTS constructs, only cells with an average acceptor intensity within a pre-specified range were analyzed. This range was set to [1000 40000] for mTFP1-Venus-based sensors or [600 24000] for Clover-mRuby2-based sensors, resulting in exclusion of <10% of cells. Finally, for VinTS constructs, cells that were not fully spread were also excluded from analysis.

### Segmentation and analysis of VinTS in FAs

Post-processing of FRET images to segment and quantify the characteristics of individual FAs was performed using custom-written code in MATLAB (Mathworks). Briefly, FAs were identified and segmented on the acceptor channel using the water algorithm (*Zamir et al., 1999*). The resultant mask was applied across all images for ease of data visualization and quantification. For each identified FA, parameters describing its brightness in the acceptor channel, morphology, and molecular loading (FRET) were determined. To identify single cells, closed boundaries were drawn by the user based on the unmasked acceptor-channel image. From these cell outlines, parameters describing cell morphology and FA subcellular location were also determined.

Line scans of single FAs were performed using ImageJ software (US National Institutes of Health, Bethesda, MD; http://imagej.nih.gov/ij/). Specifically, the line tool was used to visualize the acceptor channel intensity profile across single, large FAs in the cell periphery. The coordinates of these lines, drawn axially starting from the tip of FAs distal to the cell body, were then transferred to masked FRET efficiency images. Acceptor intensity and FRET efficiency profiles from single FAs were saved as text files for subsequent analysis.

### FRET efficiency calculations from spectrofluorometry

Hypotonic lysates were prepared from HEK293 cells as previously described (*Chen et al., 2005*). In addition to experimental samples, lysates from an equal number of untransfected cells were harvested and used as a reference background. Spectrofluorometric measurements were made with a Fluorolog-3 (FL3-22, HORIBA Scientific Jobin Yvon, Edison, NJ) spectrofluorometer with 1 nm step size, 0.2 s integration time, and 3 nm excitation and emission slit widths for all samples. For FRET measurements of mTFP1-Venus sensors, spectra were traced from 472 to 650 nm following donor excitation (λ_*Dex*_) at 458 nm, and from 520 to 650 nm following acceptor excitation (*λ*_Aex_) at 505 nm. For Clover-mRuby2 sensors, spectra were traced from 520 to 700 nm following donor excitation at 505 nm, and from 590 to 700 nm following acceptor excitation at 575 nm. The same settings were used to measure the emission spectra of full length and minimal FPs to confirm their spectral properties individually. Custom-written code in MATLAB (Mathworks) was used to calculate FRET efficiency via the (ratio)A method (*Majumdar et al., 2005*) as:

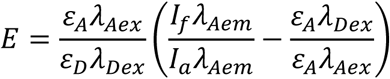

where *I_f_* and *I_a_* are the intensities, at peak acceptor emission wavelength (λ_*Aem*_, 530 nm for Venus, 610 nm for mRuby2), of the sample excited at donor and acceptor wavelengths, respectively. Donor and acceptor molar extinction coefficients (*ε_D_* and *ε_A_*, respectively) were calculated from absorbance spectra measured on the same Fluorolog-3 spectrofluorometer in absorbance mode (1 nm step size, 0.1 s integration time, 2 nm excitation and emission slit widths) using previously-measured maximal extinction coefficients: 64,000 M^−1^cm^−1^ for mTFP1 (*Ai et al., 2006*), 93,000 M^−1^cm^−1^ for Venus (*Nagai et al., 2002*), 111,000 M^−1^cm^−1^ for Clover (*Lam et al., 2012*), and 113,000 M^−1^cm^−1^ for mRuby2 (*Lam et al., 2012*). This approach was also used to confirm the absorbance spectra of the minimal FPs were unaltered as compared to the parent version.

### Statistics, bootstrapping, and data digitization

All statistical analyses, except numerical bootstrapping, were performed using JMP Pro 12 software (SAS, Cary, NC). ANOVAs were used to determine if statistically significant differences (p < 0.05) were present between groups. If statistical differences were detected, Tukey’s HSD post-hoc testing was used to perform multiple comparisons and assess statistical differences between individual groups (see **Supplementary file 3** for exact p-values and multiple comparisons test details). Box-and-whisker diagrams (Figure 1F, Figure 1–Figure supplement 2C) display the following elements: center line, median; box limits, upper and lower quartiles; whiskers, 1.5x interquartile range; red filled circle, mean; open circles, outliers.

Numerical bootstrapping using the built-in Matlab (Mathworks) function *bootstrp.m* was used to calculate 95% confidence intervals for measurements of persistence length (*L_P_*). Specifically, for each of 200 bootstrapped samples, drawn with replacement from the pertinent dataset, the *L_P_* that best reflected that sample was calculated by chi-squared error minimization. The mean and 95% confidence intervals of the resultant 200 values of *L_P_* are reported in Figure 2-Supplementary table 1. Fluorescence-force spectroscopy data (*Brenner et al., 2016*) was digitized using the digitize2.m function in Matlab (Mathworks). To recapitulate the uncertainty in these published unloaded FRET and FRET-force datasets, random sets of 100 data points obeying a gaussian distribution with the reported mean and standard deviation were used.

For FRET efficiency measurements, numerical bootstrapping of pilot data was used to determine the sample size required to estimate FRET efficiency to within 1% of the true population mean. This was determined to be 10-20 cells from 3 independent experiments for *in cellulo* measurements, or 5 independent samples for *in vitro* spectral FRET characterization. For fluorescent protein absorbance/emission spectra characterization, sample size was not pre-determined. Rather, the reproduction of data from independent experiments was deemed sufficient to draw conclusions about changes in the fluorescent protein spectral properties.

## Acknowledgements

The authors thank Dr. Ben Fabry and Dr. Wolfgang H. Goldmann (Friedrich-Alexander-Universitat Erlangen-Nurnberg) for providing vinculin -/- cells, and Dr. Harold Erickson (Duke University) and Dr. Wendy Gordon (University of Minnesota) for comments and critical examination of the manuscript.

## Author contributions

B.D.H. conceived of, designed, and supervised the study. A.S.L. and M.E.B. created and characterized expression constructs. A.S.L and A.D.L. performed FRET experiments and analyzed data. A.S.L. and M.E.B. wrote image analysis software. A.S.L., B.D.H., and A.D.L. wrote the calibration model, and A.S.L. wrote the structural model. A.S.L. and B.D.H. wrote the manuscript with input from all authors.

## Figure supplements for Figure 1

**Figure 1–Figure supplement 1.**
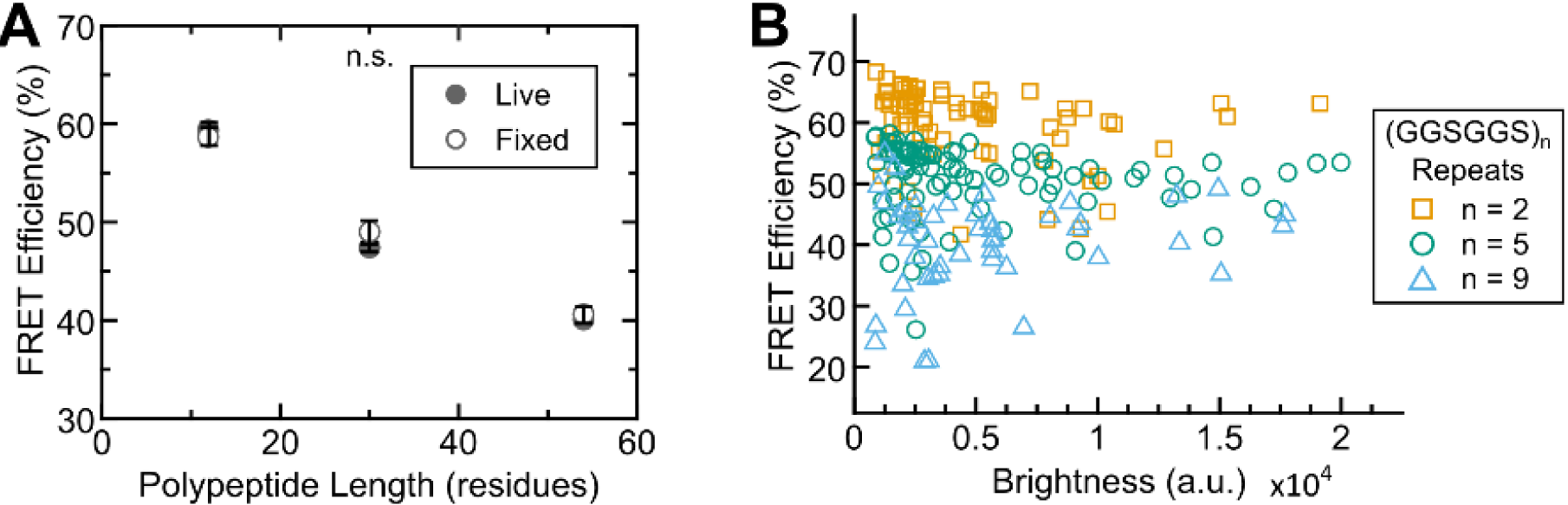
FRET efficiency measurements are insensitive to fixation and sensor intensity. (**A**) Quantification of FRET-polypeptide length relationship for “minimal” Clover-mRuby2 based TSMods *in cellulo* either live or fixed; each point represents at least n = 9 cells per experiment from three independent experiments; analysis of covariance (ANCOVA) was used to provide a model-independent assessment of statistical differences; ANCOVA interaction term p > 0.05 indicates that the relationship between FRET efficiency and polypeptide length is not significantly different (n.s.) between live and fixed conditions; error bars, s.e.m. (**B**) FRET efficiency measurements as a function of mean acceptor intensity (brightness) for fixed cells expressing TSMods consisting of the “minimal” Clover-mRuby2 FRET pair and (GGSGGS)2,5,9 extensible domains (R^2^ = 0.06, 0.01, 0.03 and n = 74, 86, 48 cells, respectively; data pooled from three independent experiments).

**Figure 1–Figure supplement 2.**
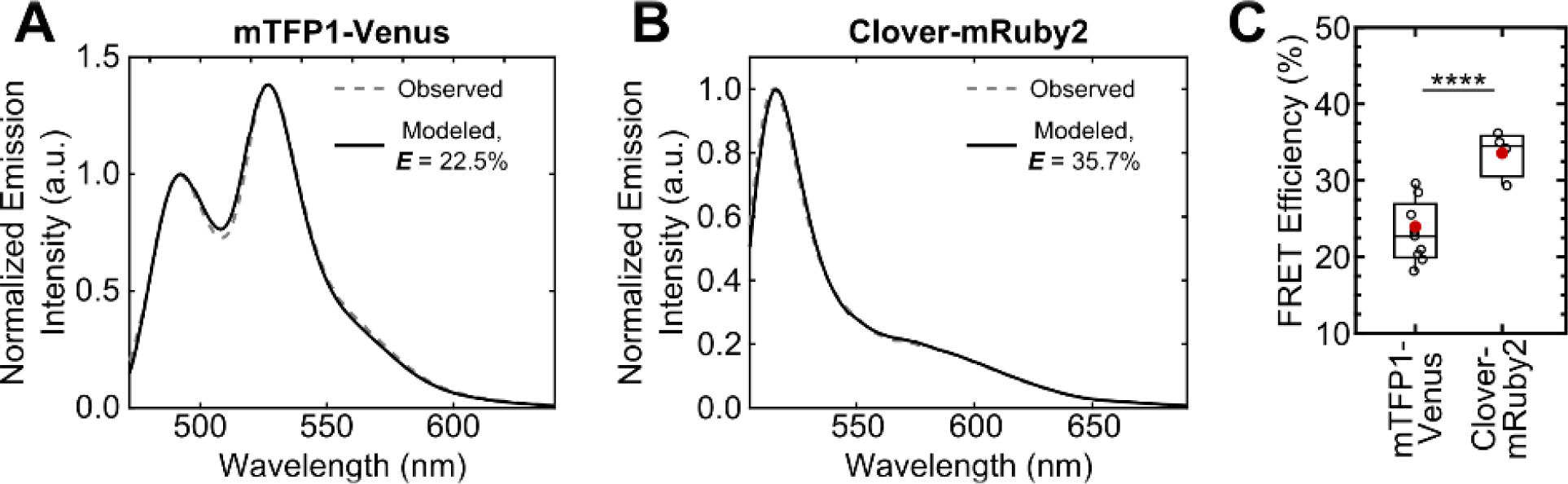
Increase in unloaded FRET efficiency with Clover-mRuby2 sensors *in vitro.* (**A, B**) Representative images of quantitative spectral analysis of mTFP1-Venus (**A**) and Clover-mRuby2 (**B**) TSMod fluorescence in cell lysates using the (ratio) A method (*Majumdar et al., 2005*). (**C**) Quantification of unloaded FRET efficiency for mTFP1-Venus and Clover-mRuby2 TSMods with (GPGGA)8 extensible domain; (n = 9 and 4 independent experiments, respectively); red filled circle denotes sample mean; *\*\*\*\** p < 0.0001, Student’s t-test, two-tailed, assuming unequal variances.

**Figure 1–Figure supplement 3.**
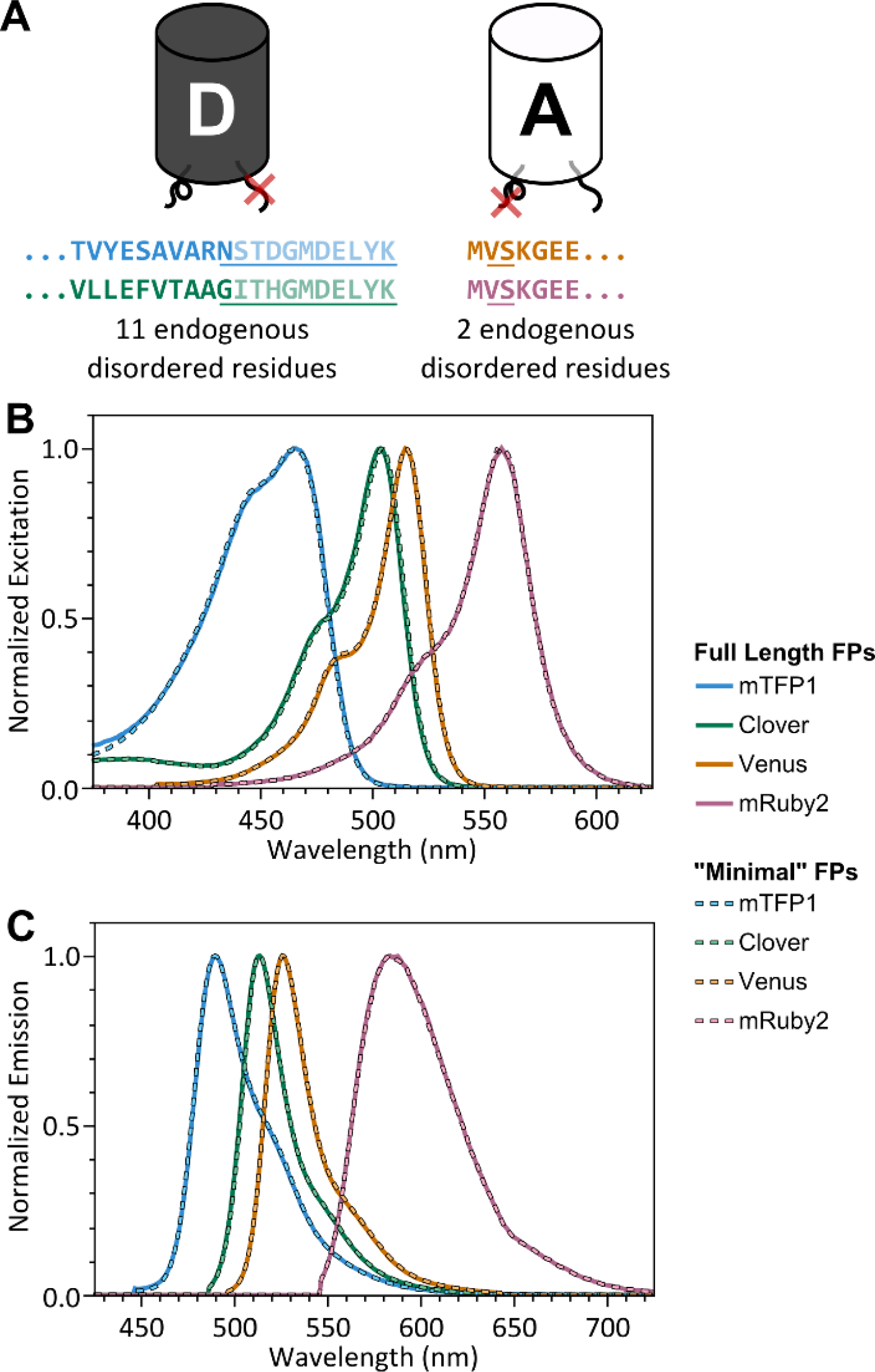
“Minimal” FPs exhibit spectral properties indistinguishable from full-length parent FPs. (**A**) Schematic of donor and acceptor FPs highlighting 11 C-terminal residues (donor FP) and 2 N-terminal residues (acceptor FP), which do not contribute to beta barrel structure, but are highly conserved between various FPs; residues not appearing in crystal structures are faded (PDB 2HQK and 1MYW for donor and acceptor FPs, respectively); residues that were removed are underlined. (**B, C**) Normalized excitation (**B**) and emission (**C**) spectra for full length and “minimal” versions of mTFP1, Venus, Clover, and mRuby2.

## Figure supplements for Figure 2

### Figure 2-Supplementary note 1: Development, validation, and implementation of a computational model describing the mechanical sensitivity of TSMods

In this supplementary note, we present a modeling-based approach to predict the relationship between applied force and measured FRET efficiency in a collection of genetically-encoded fluorescent protein (FP)-based molecular tension sensors that use unstructured polypeptides as the extensible domain (overview in Section I). This calibration model consists of a combination of accurate numerical predictions of biopolymer extension under load (Section II), simple empirical estimates correcting for the finite size (Section III) and dynamics of the FPs (Section IV), and established distance-FRET relationships (Section V). The effects of force on tension sensor module (TSMod) FRET efficiency are simulated using a Monte Carlo calculation scheme described in Section VI. To develop a modelling-based calibration approach for TSMods based on unstructured polypeptides, we investigated the ability of the proposed model to describe TSMod mechanical behaviors in both unloaded (Section VII) and loaded (Section VIII) conditions. Finally, we provide a comparison of the proposed model to other previous estimates of TSMod mechanical behavior (Section IX).

We demonstrate that the proposed model predicts the relationship between mechanical forces and FRET efficiency with sufficient accuracy to aid in the design of TSMods. Specifically, we show that the model can describe multiple experimental datasets with physically accurate parameters (Figure 2–Supplementary table 2). Furthermore, calibrations suggest that vinculin tension sensors based on these TSMods report forces that are consistent with estimates from previously developed tension sensors (*Grashoff et al., 2010*) as well as those from traction force microscopy (*Balaban et al., 2001*).

### I. Overview of model

Estimates of TSMod mechanical sensitivity relate measured FRET efficiencies to applied forces. However, force and FRET are neither linearly nor simply related, as the relationship will depend on the mechanical properties of the extensible domain as well as the manner in which the extensible domain is connected to the fluorescent moiety. When considering TSMods with unstructured polypeptide extensible domains, there are two key distance metrics that must be considered. First, the end-to-end distance of the polypeptide (*r*_e_) can be directly related to force via various mechanical models of polymer mechanics (Section II, Figure 2–Figure supplement 1A). Second, the chromophore separationdistance (*r_c_*) is directly related to FRET via established distance-FRET relationships including the Förster equation (Section IV, Figure 2–Figure supplement 1C). The relationship between *r_e_* and *r_c_* therefore is of critical importance and requires knowledge of the radii of the FPs (Section III, Figure 2–Figure supplement 1B. Details of how the calculations were performed are provided in Section V.

**Figure 2–Figure supplement 1.**
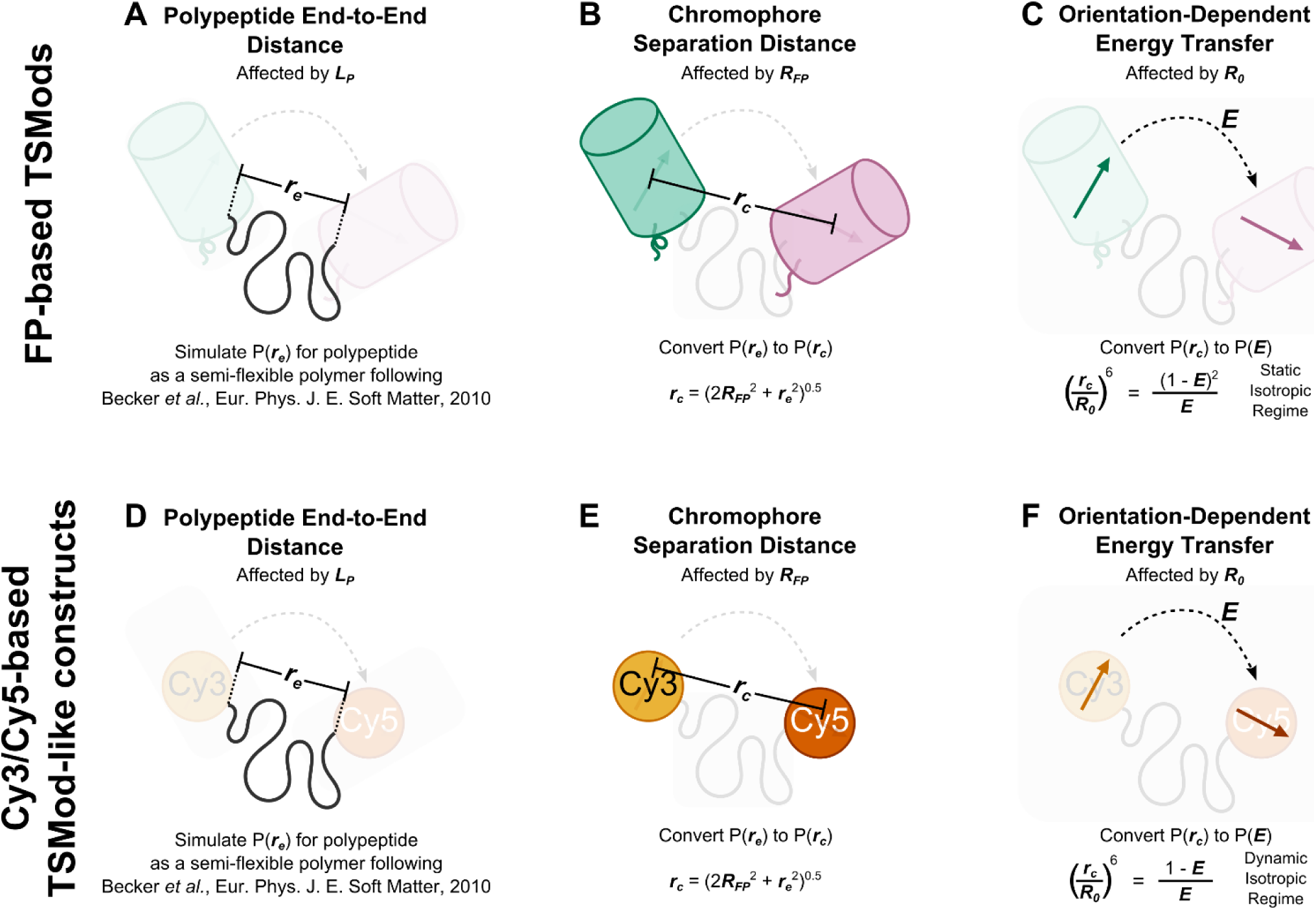
Schematic depiction of biophysical model describing the mechanical sensitivity of TSMods. (**A-C**) Mechanical sensitivity of FRET-based TSMods depends on the mechanical properties and length of the extensible domain (**A**), the physical separation of the chromophores within the FPs (**B**), and the relative orientation and timescale of rotation of the FPs (**C**). (**D-F**) Similarly, the mechanical sensitivity of FRET-based TSMod-like constructs which utilize Cy3 and Cy5 as the fluorescent moiety depends on the mechanical properties and the length of the extensible domain (**D**), the physical separation of the organic dyes (**E**), and the relative orientation and timescale of rotation of the dyes (**F**). Note when modeling large FP-based TSMods, the static isotropic regime of energy transfer is valid, while FRET between organic dyes is more appropriately described by the dynamic isotropic assumption of the classical Förster equation (*Vogel et al., 2014*).

### II. Extensible domain mechanics are well-described by the worm-like chain model

To describe the mechanics of the unstructured polypeptides that serve as the extensible elements of the TSMods, we employ the worm-like chain (WLC) model. This model, widely used in the study of the mechanics of biopolymers, captures the mechanical characteristics of a polymer in two key length scales, the persistence length (*L_P_*) and the contour length (*L_c_*). The persistence length describes the stiffness of the polymer, and is defined as the length over which the tangent direction of the polymer remains correlated (*Marko and Siggia, 1995*). The contour length describes the fully extended length of the polymer, and is defined by the product of the monomer size (*b_0_* = 0.38 nm for amino acid chains (*Evers et al., 2006a*)) and the number of monomers in the chain (*N*). Most often, the WLC model is presented as an approximation of the predicted force-extension curve, relating the average extension of the polymer to an applied force (*Marko and Siggia, 1995*):

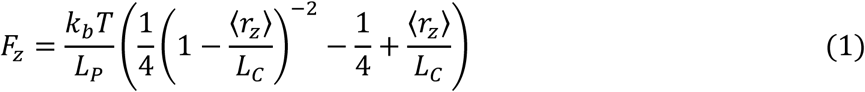

where *F_z_* is the force on the polymer chain, 〈*r_z_*〉 is the average chain extension, *k_b_* is the Boltzmann constant, and *T* is temperature. A more accurate force-extension relation was developed by (*Bouchiat et al., 1999*) by a simple 7^th^ order polynomial correction scheme:

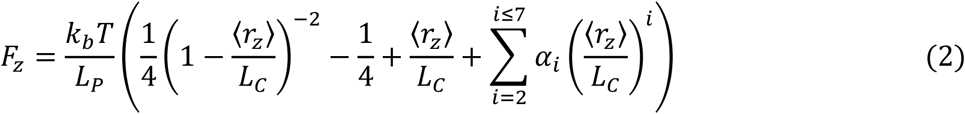

where *α*_2_ = –0.5164228, *β*_3_ = –2.737418, *α*_4_ = 16.07497, *α*_5_ = –38.87607, *α*_6_ = 39.49944, and *α*_7_ = –14.17718.

However, despite the widespread and successful use of these approximations of the force-extension curve of a WLC in the quantification of biopolymer mechanics (see Figure 2-Supplementary table 2), these relationships are not directly compatible with most FRET-based measurements for several reasons. The issue most pertinent to this work is that Eq. 1 and 2 relate force to the average extension (〈*r_z_*〉) of the polypeptide chain, which is zero in the absence of force. However, the non-zero polypeptide end-to-end distance (*r_e_*), which describes the finite rest length of the extensible domain, is more directly related to the chromophore separation distance (*r_c_*), which is the key distance determining FRET efficiency (E). Additionally, the nonlinear relationships between these quantities (*r_e_*, *r_c_*, and *E*) prevent the use of simple heuristic correction schemes to relate various distance metrics (*Evers et al., 2006a*).

Previous work has developed formalisms that enable these concerns to be accounted for (*Becker et al., 2010; Evers et al., 2006a; Vogel et al., 2012; Vogel et al., 2014*), but they have yet to be applied to the development of FRET-based TSMods. A key realization is that TSMods are molecular-scale tools and therefore will be subject to thermal fluctuations (*Vogel et al., 2012*). Therefore, to begin it is necessary to describe the end-to-end distance of the extensible polypeptide (r_e_) as a probability density function P(r_e_), which is obtained by integrating the end-to-end vector probability density function 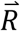 over all the angles through which the chain may rotate:

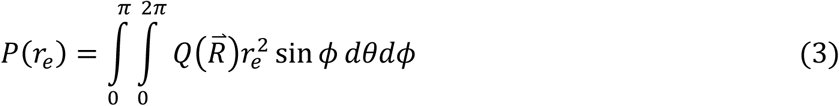

yielding

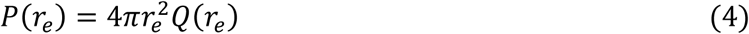

Forces will alter this distribution following the widely-applied Boltzmann relation:

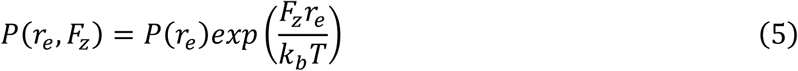

where *P(r_e_,F_z_*) is the probability of observing a given end-to-end distance for the polymer chain at a given force *F_z_*. The average extension of the polypeptide chain (〈r_z_〉), which is similarly affected by forces following the Boltzmann relation, was calculated as:

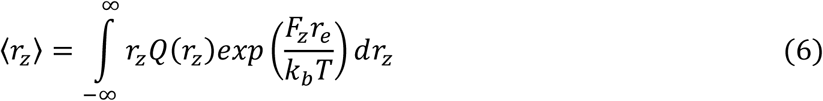

However, as stated above, this parameter has limited relevance for further FRET calculations and was only calculated for comparison to classical WLC force-extension relationships (Eq. 1 & 2).

A variety of closed-form equations that describe the behavior of biopolymers exist, but are accurate only in specific force regimes and ranges of polymer mechanics (*Becker et al., 2010*). For example, the second Daniels approximation (*Daniels, 1952; Yamakawa and Stockmayer, 1972*) describes *P(r_e_*) for small extensions of soft polymers (*L_C_/L_P_* > 5) following:

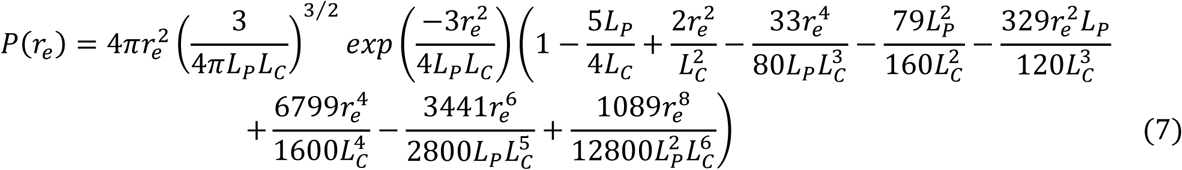

In other work Wilhelm & Frey develop an approximation that describes the behavior of short chain or stiff (*L_C_/L_P_ < 2*) polymers (*Wilhelm and Frey, 1996*). Specifically Wilhelm & Frey provide two series expansions:

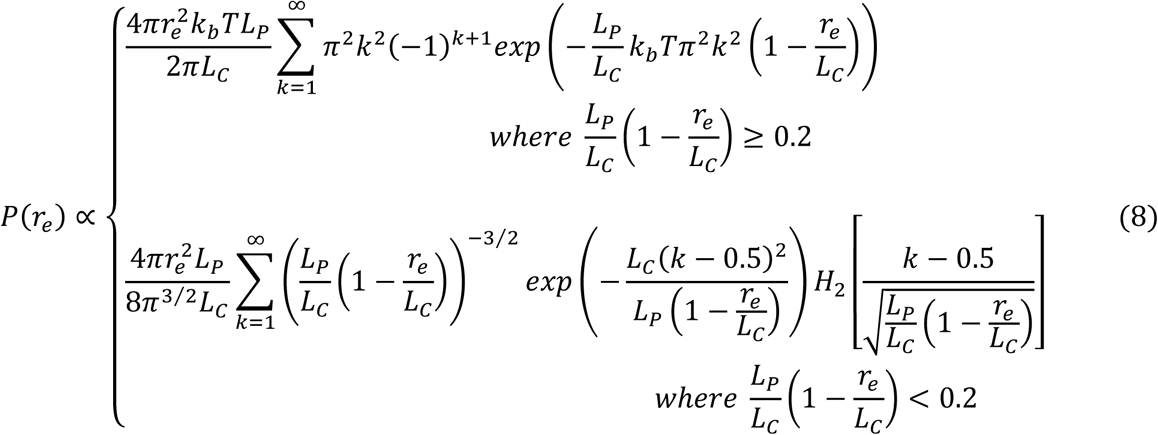

Where *H_2_* is the second Hermite polynomial *H_2_* = 4*x*^2^ – 2. However, these approximations were later found to be valid primarily at high extension (*Becker et al., 2010*).

Excitingly, Becker *et al.* developed an explicit expression which interpolates between a number of these approximations, including but not limited to those described above, and created an ansatz that accurately describes *P(r_e_*) over a wide range of polymer mechanics 0.05 < *L_C_/L_P_* < 50 (*Becker et al., 2010*), and likely to much higher values in the absence of excluded volume effects. Normalizing the stiffness to the contour length such that *K = L_P_/L*_C_, the final form of the interpolated probability density function *P〈r_e_〉* is given by:

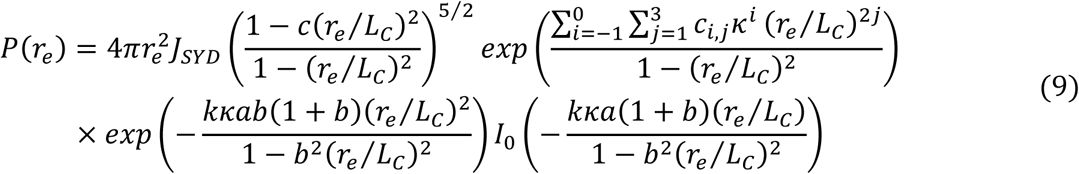

where *J_SYD_* is the Shimada-Yamakawa J-factor:

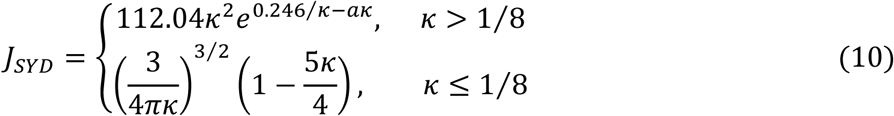

with constants *a = 14.054, b = 0.473,* and coefficients *Ci,j* given as:

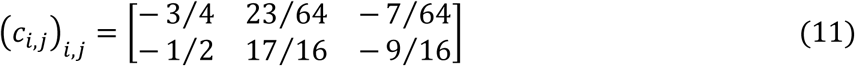

Additionally, *I*_0_ is a modified Bessel function of the first kind and parameters *c* and *d* are defined as:

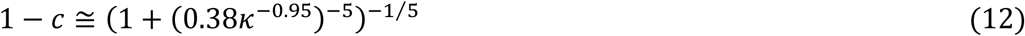

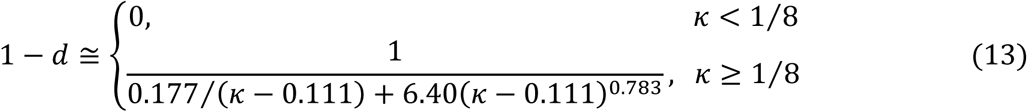

To describe the behavior of the unstructured polypeptides used in the assembly of TSMod constructs, values of *L_c_*, *L_P_* and *F_z_* were specified, and the inverse-transform-sampling method (*Titantah et al., 1999*) was used to generate sets of random numbers distributed according to *P(r_e_,F_z_*) as specified by combining the Becker *et al.* ansatz (Eq. 9) with the Boltzmann relation (Eq. 5). Note that while this describes the mechanical behavior of the unstructured polypeptide, the overall force sensitivity and FRET output from a given TSMod construct will also depend on the radii and photophysical properties of the FPs attached to this extensible domain as well (discussed further in Sections III and IV).

To confirm that this formalism was both correctly implemented and suitable for the intended purposes, several calculations demonstrating the ability of our simulations to accurately reproduce previous results were completed. First, we verified the ability of the model to described the unloaded states of an unstructured polypeptide by comparing the zero-force state of the calculated *P(r_e_*) (Eq. 9) to published simulations of a discretized version of the WLC model developed by Wilhelm & Frey that is often used for stiffer polymers (*Wilhelm and Frey, 1996*), and the second Daniels approximation for softer polymers (*Daniels, 1952; Yamakawa and Stockmayer, 1972*), observing excellent agreement in all cases (Figure 2–Figure supplement 2A). Next, we determined the predictions of the scaling between the average end-to-end distance (〈r_e_〉) and polypeptide length (*N*) for the proposed model, observing the classical 〈r_e_〉∼*N*^0.5^ behavior characteristic of random chains (Figure 2–Figure supplement 2B). Finally, the model was used to generate force-extension curves (Eq. 6) for various ratios of contour length to persistence length and compared to the most accurate numerical approximation of the force-extension curve of a WLC (*Bouchiat et al., 1999*) (Eq. 2). The model agrees well with the numerical approximation across all *L_C_/L_P_* ratios (Figure 2–Figure supplement 2C). Together these data show that this approach accurately simulates *P(r_e_,F_z_*) for the expected mechanics (Figure 2-Supplementary table 2) and lengths achievable for the extensible domain of polypeptide-based TSMods.

### III. Effects of fluorescent proteins

Next, we sought a method to account for the finite size of the FPs attached to the ends of the extensible domain. Like any FRET-based biosensor, the pertinent energy transfer distance for a TSMod is the inter-chromophore distance (*r_c_*), not simply *r_e_*. Since the FPs have finite size, the difference between these two quantities is not negligible and has been approximated by several methods (*Brenner et al., 2016; Evers et al., 2006a*). In light of the experimental complexities associated with single molecule fluorescence force spectroscopy, Brenner *et al.* utilized a simple approach and approximated *r_c_* =*r_e_* + C (*Brenner et al., 2016*). However, it is generally recognized that FPs are free to rotate around the ends of the polypeptide in a random fashion requiring more complex relationships to relate *r_c_* to *r_e_*. Attempts by Evers *et al.* to recapitulate this behavior by simulating 66 possible orientations for each FP at a given *r_e_* (4356 total conformations possible) and considering the average as an appropriate estimate of *r_c_* are a reasonable means of estimating these effects (*Evers et al., 2006a*). However, Evers’ approach does not yield a realistic limit at short polypeptide lengths. Specifically, as the polypeptide length approaches zero, this approach allows for the centers of the FPs to be closer than the sum of their individual radii (Figure 2–Figure supplement 2D). To address this issue and approximate the effects of steric hindrance between the two FPs, we used a heuristic approach involving a quadrature sum of the key distances in this problem:

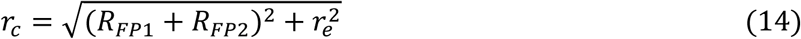

where *R_FP1_* and *R_FP2_* are the radii of the donor and acceptor FPs, respectively. Based on crystal structures of FPs (*Ai et al., 2006; Ormo et al., 1996; Rekas et al., 2002*) and estimates of FP hydrodynamic radii from fluorescence correlation spectroscopy (*Hink et al., 2000*), we set *R_FP1_ = R_FP2_* = 2.3 nm. For cyanine dyes, these quantities were set to *R_FP1_* = *R_FP2_* = 0.95 nm, which is mathematically equivalent to the reported hydrodynamic radii of Cy3 and Cy5, which are 0.90 nm (*Muddana et al., 2009*) and 1.00 nm (*Widengren and Schwille, 2000*), respectively. This steric hindrance approximation recapitulates the behavior of Evers’ spherical integration approach for longer polypeptides while also providing a realistic limit for very short polypeptides (Figure 2–Figure supplement 2D). As the operation is always the same for a given separation distance and size of FPs, a transfer function (Eq. 14) was written to a file and treated as a lookup table for the conversion of *r_e_* to *r*_c_.

### IV. Accounting for rotational entropy phenomenologically

When applying load to a molecule, a portion of energy works against rotational entropy before the molecule is extended. This phenomenon is known to limit resolution in force spectroscopy measurements (*Neuman and Nagy, 2008*), and was therefore considered in our modeling of force sensitive biosensors. Previous descriptions of this phenomenon (*Brenner et al., 2016*) have defined a critical force:

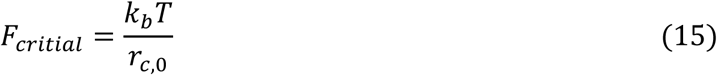

which is required to orient the molecule in the direction of pulling, and is set by the thermal energy of the system and the unloaded distance between the FP chromophores (r_c0_). Thus, forces below *F_critical_* do not entirely act to increase the separation distance between the FPs. We heuristically account for these effects using:

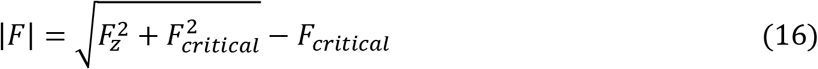

This results in only a fraction of the total amount of force applied to the TSMod, |*F*|, leading to extension of the TSMod through the force impinging on the extensible domain itself *F_z_* (Figure 2–Figure supplement 2E). This causes the emergence of a regime of relative insensitivity of the sensors at very small forces (Figure 2–Figure supplement 2F). Notably, this simple equation recapitulates observations in single molecule force spectroscopy experiments (*Brenner et al., 2016; Grashoff et al., 2010*).

**Figure 2–Figure supplement 2.**
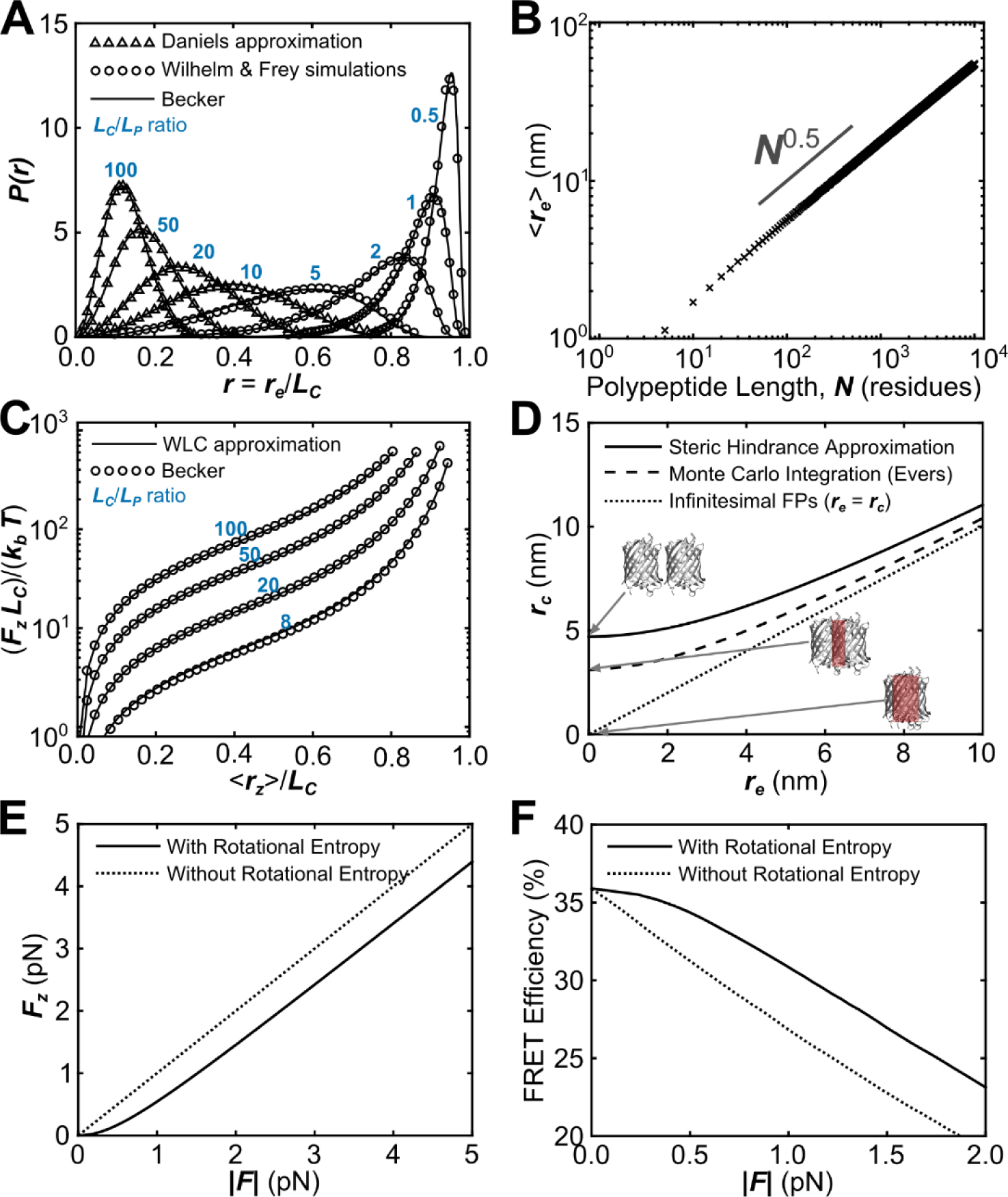
Verification of the proper implementation of a biophysical model describing the mechanical sensitivity of FRET-based TSMods. (**A**) Probability distributions of polypeptide end-to-end distance (r_e_), normalized to contour length (*L_C_*) such that *r = r_e_/L_c_* calculated using the second Daniels approximation (*Daniels, 1952; Yamakawa and Stockmayer, 1972*), simulations from (*Wilhelm and Frey, 1996*), and an ansatz approximation used in the proposed model based on (*Becker et al., 2010*) that smoothly bridges between various mechanical regimes. (**B**) Demonstration that the numerical ansatz predicts the standard length 〈*r_e_〈∼N^1/2^* scaling behavior characteristic of unstructured polypeptides. (**C**) Under load, the average extension (〈r_z_〉) of the polypeptide extensible domain predicted by the (*Becker et al., 2010*) ansatz closely follows the previous prediction of the force-extension curve for a worm-like chain (*Bouchiat et al., 1999*). (**D**) Possible approaches to estimate the chromophore separation distance (r_c_), from polypeptide end-to-end distance 〈r_e_〉. Note that some approaches predict unphysical overlap of the FPs at short linker lengths, which is highlighted in red for the extreme condition of infinitesimal linker length. (**E, F**) The proposed model includes a dimensionless group (determined by the applied load, the rest length of the construct, and thermal energy) that heuristically accounts for the effects of rotational entropy. This results in a reduced amount of force transferred through FPs to the polypeptide at low applied force (**E**); this difference between the force causing extension of the polypeptide (F_z_ ) and applied force (|*F*|) manifests in a plateau in the FRET-force relationship at very small loads (**F**).

### V. Förster resonance energy transfer

FRET efficiency (*E*) was calculated for each *r_c_* in the set using either the Förster equation:

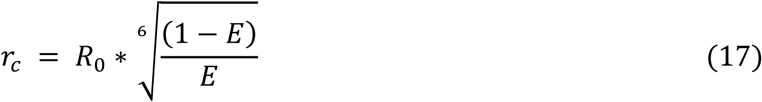

which assumes a dynamic isotropic regime for fluorescent moiety rotation (*κ*^2^ = 2/3) and is applicable to only quickly rotating, small fluorescent moieties (e.g. cyanine dyes), or an alternative distance-FRET relationship:

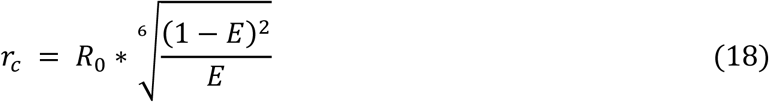

which is valid in the static isotropic regime and more accurately describes energy transfer between fluorescent moieties (e.g. FPs) which rotate on timescales longer than those associated with FRET (*Vogel et al., 2012; Vogel et al., 2014*). The Förster radius (*R_0_*) appears in both relationships and is defined as the distance at which a pair of fluorescent moieties exhibit 50% FRET efficiency. The value for *R*_0_ was set, based on previous measurements, to 5.7 nm for the mTFP1-Venus FRET pair (*Ai et al., 2006*), 6.3 nm for the Clover-mRuby2 FRET pair (*Lam et al., 2012*), and 5.4 nm for the Cy3-Cy5 FRET pair (*Buckhout-White et al., 2014; Sanborn et al., 2007*). As before, either transfer function was written to a file and treated as a lookup table for the conversion of any *r_c_* to *E.* The mean of the resultant FRET distribution was then reported as the FRET efficiency at a given force.

### VI. Calculations

Monte Carlo simulations were used to determine the average polypeptide extension (〈*r_z_*〉) as well as probability distributions for the polypeptide end-to-end distance (*r_e_*), FP chromophore separation distance (*r_c_*), and FRET efficiency (*E*) at a given force on the polypeptide (*F_z_*), persistence length (*L_P_*), and number of residues in the polypeptide (*N,* used to calculate contour length, *L_C_ = N * b_0_*). First, a set of at least 10,000 random numbers obeying the probability distribution function *P(r_e_*) described by the Becker *et al.* ansatz (Eq. 9) was generated from a uniform distribution using the inverse-transform sampling method (*Titantah et al., 1999*). Then, the impact of force on *P(r_e_*) was accounted for following the Boltzmann relation (Eq. 5). Next, *P(r_e_*) was first converted to *P(r_c_*) using the *r_e_*-to-r_c_ lookup table described by Eq. 14, and then *P(E*) using the *r_c_-to-E* lookup table described by Eq. 17 for TSMod-like constructs with cyanine dyes as the fluorescent moiety or Eq. 18 for FP-based TSMods. Additionally, the force affecting the extension of the sensor (|*F*|) was determined from *F_z_* following Eq. 16. Finally, the average of each population distribution as well as |*F*|, *N,* and *L_P_* were written to a line of a TSMod calibration text file. These operations were repeated from 0 to 50 pN of force at increments of 0.1 pN to map out the FRET efficiency (*E*) as a function of force (|*F*|) and extension (*r_z_*) with a given set of parameters.

### VII. Model validation in unloaded conditions

To evaluate whether the model can be used to describe the behavior of TSMods in unloaded conditions a variety of experimental results were evaluated. First, we investigated whether the model accurately describes the relationship between FRET efficiency and polypeptide length for Clover-mRuby2 TSMods and provide reasonable estimates of *L_P_* for (GGSGGS)_n_ or (GPGGA)_n_ extensible domains in both *in vitro* and *in cellulo* environments. To limit the parameter space, estimates for the other two input parameters (*R*_0_ and *R_FP_*) were obtained from previous reports of the Förster radius of Clover and mRuby2 (*R_0_ = 6.3* nm (*Lam et al., 2012*)) and the hydrodynamic radii of GFP family FPs (*R_FP_ = 2.3* nm (*Hink et al., 2000*)). This leaves *L_P_* as the sole adjustable parameter. Simulations were then used to predict the relationship between FRET efficiency and polypeptide length for a variety of polypeptide chains (10 *< *N* < 100 residues) and persistence lengths (0.1 < *L_P_* < 2.0* nm, 0.01 nm increments) and compared to *in vitro* FRET efficiency measurements of TSMods in cell lysates (Figure 1G). Model fits and bootstrapped 95% confidence intervals indicate a quantitative difference in the mechanics of (GPGGA)_n_ (*L_P_ = 0.74* ± 0.05 nm) and (GGSGGS)_n_ (*L_P_* = 0.33 ± 0.05 nm) extensible domains *in vitro.* Due to the presence of proline residues in the repeated structure of the (GPGGA)_n_ extensible domain, this stiffer mechanical response is not unexpected (*Becker et al., 2003*). In contrast, fits to sensitized emission-based FRET efficiency measurements of sensors expressed *in cellulo* revealed quantitatively indistinguishable FRET-polypeptide length relationships for TSMods containing (GPGGA)_n_ (*L_P_* = 0.50 ± *0.02* nm) and (GGSGGS)_n_ (*L_P_* = 0.48 ± 0.05 nm) extensible domains (Figure 1H). These values agree well with previous reports of both (GPGGA)_n_ and (GGSGGS)_n_ mechanics (*Becker et al., 2003; Evers et al., 2006a*). In total, these results confirm that there is a quantitative difference in extensible domain mechanics between *in vitro* and *in cellulo* environments, although less-so for (GGSGGS)_n_-based TSMods, and provide quantitative measurements of *L_P_* that can be used to generate *in cellulo* TSMod calibrations.

To rule out the possibility that constraining *R_FP_* and/or *R*_0_ to their published values was skewing estimates of *L_P_*, we examined whether improved fits to experimental data could be obtained if either *R_FP_* or *R*_0_ (in addition to *L_P_*) were left unconstrained. Specifically, we compared simulated FRET-polypeptide length relationships for various combinations of either *L_P_* and *R*_0_ (at *R_FP_ = 2.3* nm) or *L_P_* and *R_FP_* (at *R*_0_ = 6.3 nm) to experimental FRET-length measurements made *in vitro* (Figure 2–Figure supplement 3) or *in cellulo* (Figure 2–Figure supplement 4). Simulations included *L_P_* ranging from 0.10 to 1.00 nm, *R_FP_* from 1.7 to 3.4 nm, and *R*_0_ from 5.1 to 6.8 nm. We then evaluated the chi-squared error between model predictions and experimental data for each such combination of *L_P_* and *R_FP_* (Figure 2–Figure supplement 3A, 3B, 4A, 4B) or *L_P_* and *R*_0_ (Figure 2–Figure supplement 3C, 3D, 4C, 4D). In all cases, regardless of measurements made *in vitro* or *in cellulo,* minimal chi-squared error between model and experiment occurs at or close to the literature estimates of both *R_FP_* and *R_0_*(highlighted with vertical green rectangles). Finally, compared to the single unconstrained parameter *L_P_*, leaving two parameters unconstrained leads to no significant improvement in fits to the relationship between FRET efficiency and polypeptide length in unloaded conditions (Figure 2–Figure supplement 3E, 4E). Thus, we conclude the use of literature-based estimates for key parameters did not bias our analysis.

**Figure 2–Figure supplement 3.**
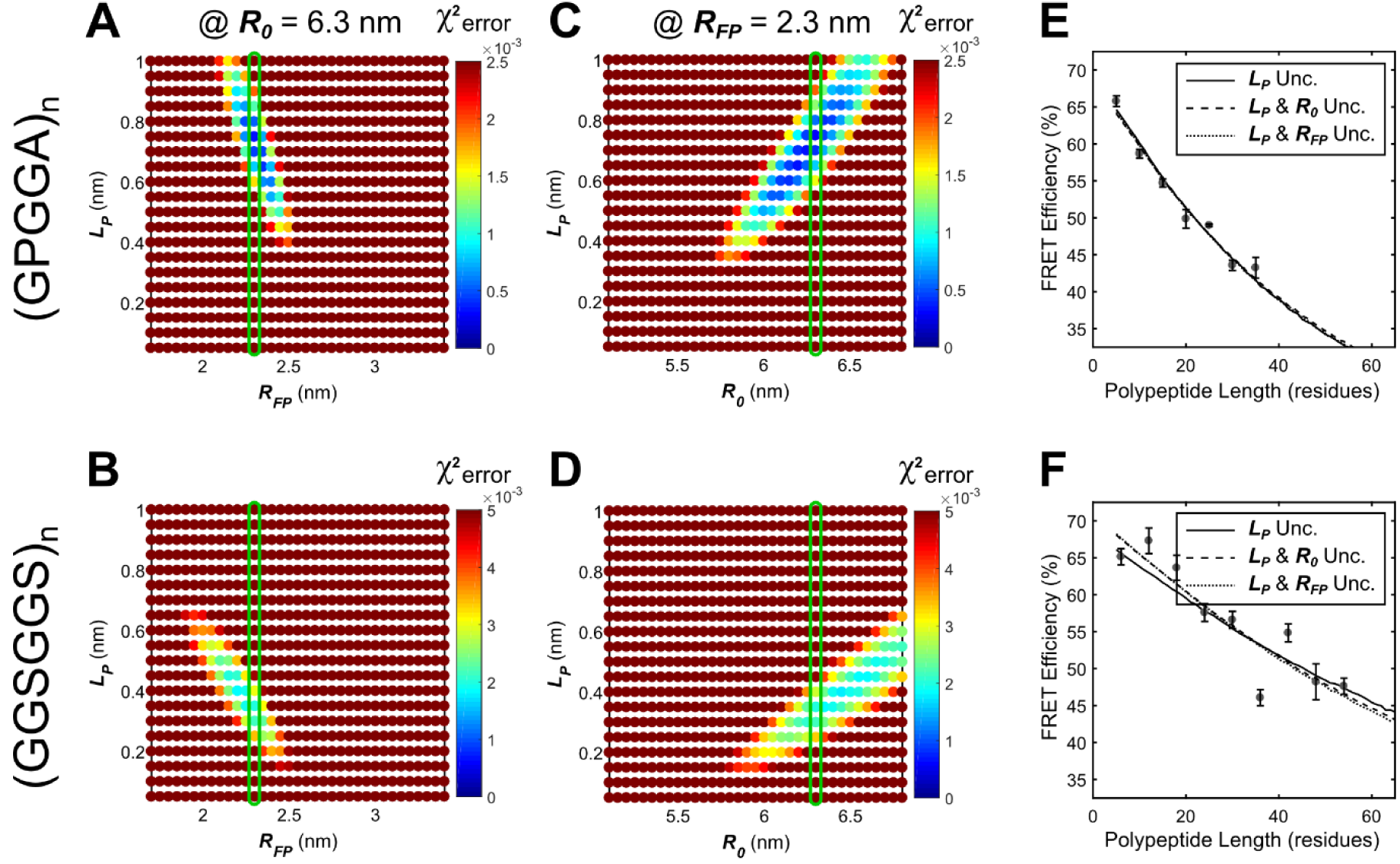
Parameter constraint has minimal effects on measurement of polypeptide persistence length (*L_P_) in vitro*.(**A**, **B**) Heatmaps of chi-squared error of model fits to *in vitro* “minimal” Clover-mRuby2 based TSMod FRET-length measurements for various *R_FP_* and *L_P_* for (GPGGA)_n_ (**A**) and (GGSGGS)_n_ (**B**) extensible domains; literature estimate of *R_FP_* used for parameter constraint highlighted with green bar; *R*_0_ is constrained to 6.3 nm. (**C**, **D**) As in panels **A** and **B**, except various *R*_0_ and *L_P_* parameters were examined; green bar highlights *R*_0_ parameter estimate; *R_FP_* is constrained to 2.3 nm. (**E**, **F**) Model fits to *in vitro* FRET-length relationships for “minimal” Clover-mRuby2 based TSMods containing the (GPGGA)_n_ (**E**) or (GGSGGS)_n_ (**F**) extensible domain are equally accurate with either a single unconstrained (unc.) parameter (*L_P_*) or two unconstrained parameters. **See Figure 2-Supplementary table 1** for full list of fit parameters compared to values from the literature.

**Figure 2–Figure supplement 4.**
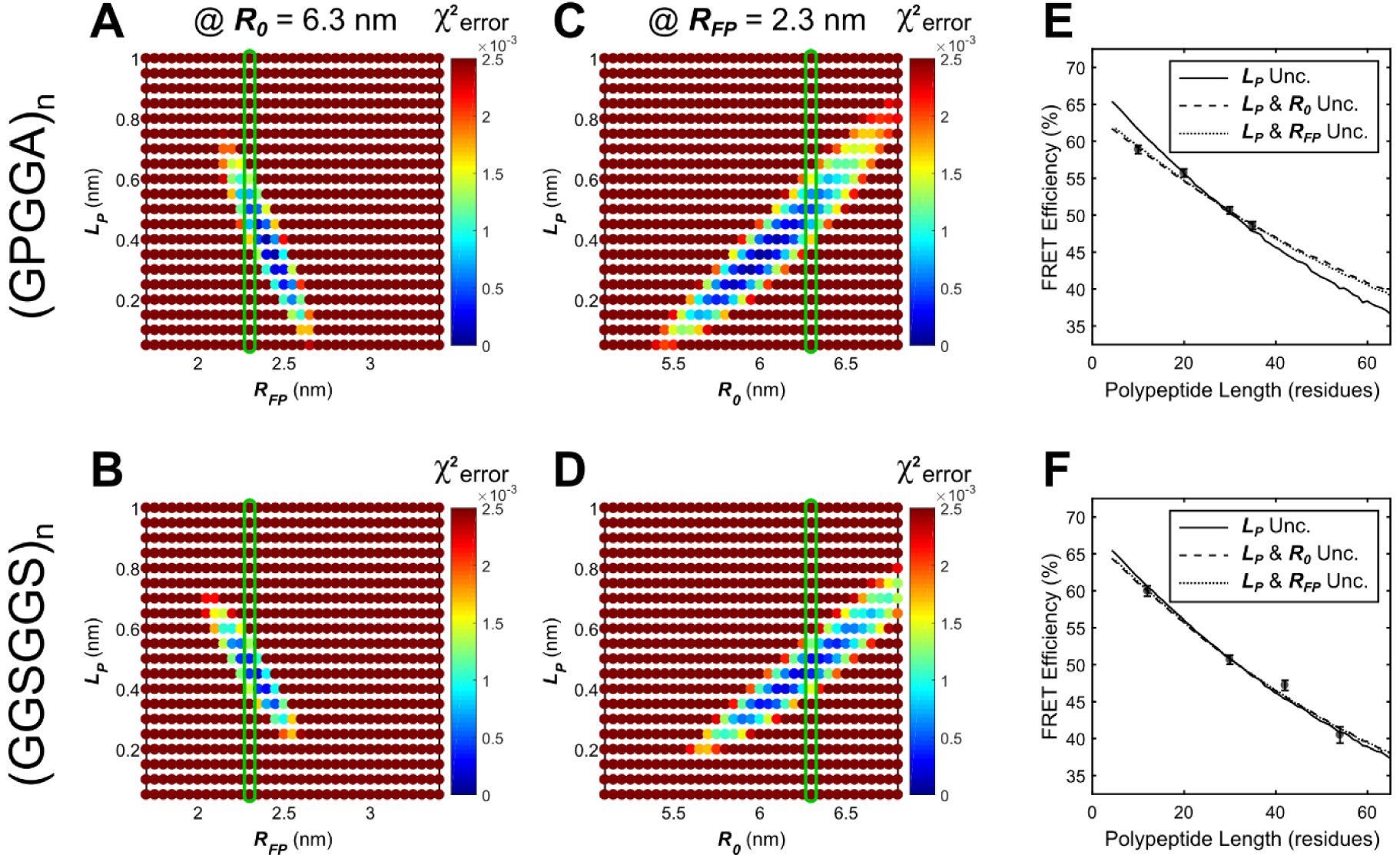
Parameter constraint has minimal effects on measurement of polypeptide persistence length (*L_P_) in cellulo.* (**A**, **B**) Heatmaps of chi-squared error of model fits to *in cellulo* “minimal” Clover-mRuby2 based TSMod FRET-length measurements for various *R_FP_* and *L_P_* for (GPGGA)_n_ (**A**) and (GGSGGS)_n_ (**B**) extensible domains; literature estimate of *R_FP_* used for parameter constraint highlighted with green bar; *R*_0_ is constrained to 6.3 nm. (**C**, **D**) As in panels **A** and **B**, except various *R*_0_ and *L_P_* parameters were examined; green bar highlights *R*_0_ parameter estimate; *R_FP_* is constrained to 2.3 nm. (**E**, **F**) Model fits to *in cellulo* FRET-length relationships for “minimal” Clover-mRuby2 based TSMods containing the (GPGGA)_n_ (**E**) or (GGSGGS)_n_ (**F**) extensible domain are equally accurate with either a single unconstrained (unc.) parameter (*L_P_*) or two unconstrained parameters. See Figure 2-Supplementary table 1 for full list of fit parameters compared to values from the literature.

### VIII. Model validation in loaded conditions

To evaluate the ability of the calibration model to describe the mechanical sensitivity of TSMods subject to mechanical loads, we examined model fits to previously published fluorescence-force spectroscopy data of (GPGGA)_n_ polypeptides labelled with Cy3 and Cy5 dyes (*Brenner et al., 2016*). These TSMod-like constructs differ from the FP-based TSMods in terms of the physical size and the Förster radius of the fluorescent moiety. Therefore, we set *r_FP1_* = 0.9 nm and *r_FP2_* = 1.0 nm based on measurements of the hydrodynamic radii of Cy3 (*Muddana et al., 2009*) and Cy5 (*Widengren and Schwille, 2000*) dyes, respectively. We also set *R*_0_ = 5.4 nm based on the reported photophysical properties of the Cy3-Cy5 FRET pair (*Buckhout-White et al., 2014; Sanborn et al., 2007*). As above, the one unconstrained parameter remaining, persistence length (*L*_P_), describes the mechanical response of the extensible domain. Simulations of Cy3-Cy5 constructs with various polypeptide lengths (10 < N < 100 residues) and persistence lengths (0.1 <L_P_ <2.0 nm) were then compared to experimentally measured FRET- polypeptide length and FRET-force relationships (Figure 2). The simulations where *L_P_∼1.05* nm agree well with both the FRET-polypeptide length relationship over the range of polypeptide lengths assessed (Figure 2A), as well as the measured relationship between FRET efficiency and force (Figure 2B). For comparison, additional FRET-polypeptide length and FRET-force relationships for a range of *L_P_* values between 1.0 nm and 1.15 nm are shown (lines in Figure 2A, shaded region in Figure 2B). Together, these data indicate that the proposed model can describe published fluorescence force measurements of TSMod-like constructs in both unloaded and loaded conditions with identical parameters.

The relatively stiff mechanical response of the (GPGGA)_n_ extensible domain (*L_P_∼1.0* nm) is somewhat unexpected given reports of other unstructured polypeptide mechanics (Figure 2-Supplementary table 2) and our own measurements of extensible domain mechanics *in vitro* (Figure 1G, Figure 2-Supplementary table 1). To rule out the possibility that constraining *R_FP_* and/or *R*_0_ to their published values was skewing estimates of *L_P_*, we examined whether improved fits to experimental data could be obtained if either *R_FP_* or *R*_0_ (in addition to *L_P_*) were left unconstrained. Specifically, we compared simulated FRET-polypeptide length and FRET-force relationships for various combinations of either *L_P_* and *R*_0_ (at 〈*R_FP_〉* = 0.95 nm) or *L_P_* and *R_FP_* (at *R*_0_ = 5.4 nm) to experimental FRET-length and FRET-force measurements (Figure 2–Figure supplement 5). Simulations included *L_P_* ranging from 0.20 to 1.25 nm, *R_FP_* from 0.3 to 2.0 nm, and *R*_0_ from 4.8 to 6.5 nm. We then evaluated the chi-squared error between model predictions and experimental data for each such combination of *L_P_* and *R_FP_* (Figure 2–Figure supplement 5A, B) or *L_P_* and *R*_0_ (Figure 2–Figure supplement 5C, D). Minimal chi-squared error between model and experiment occurs at or close to the literature estimates of both *R_FP_* and *R_0_* (highlighted with vertical green rectangles). Finally, compared to the single unconstrained parameter *L_P_*, leaving two parameters unconstrained leads to no significant improvement in fits to either the unloaded

**Figure 2–Figure supplement 5.**
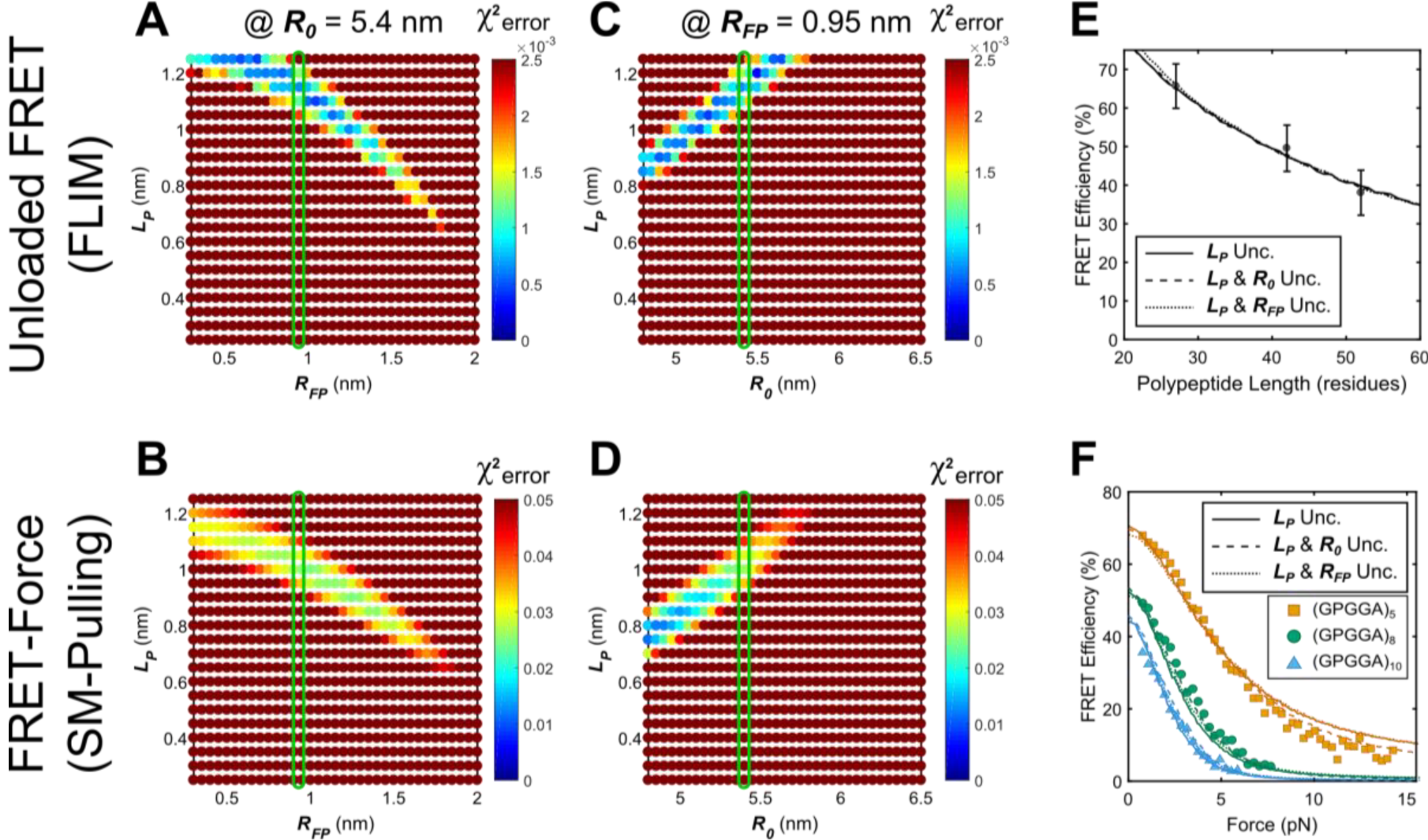
Parameter constraint has minimal effects on measurement of polypeptide persistence length (*L_P_*) for TSMod-like constructs in unloaded or loaded conditions. (**A**, **B**) Heatmaps of chi-squared error of model fits to published fluorescence-force spectroscopy measurements of (GPGGA)5,8,10 extensible domains flanked by Cy3 and Cy5 fluorescent dyes in unloaded (**A**) and loaded (**B**) conditions; literature estimate of *R_FP_* used for parameter constraint highlighted with green bar; *R*_0_ is constrained to 5.4 nm. (**C, D**) As in panels **A** and **B**, except various *R*_0_ and *L_P_* parameters were examined; green bar highlights *R*_0_ parameter estimate; *R_FP_* is constrained to 0.95 nm. (**E, F**) Model fits to published fluorescence-force spectroscopy data of TSMod-like constructs in unloaded (**E**) and loaded (**F**) conditions are equally accurate with either a single unconstrained (unc.) parameter (*L_P_*) or two unconstrained parameters. See **Figure 2-Supplementary table 1** for a full list of fit parameters compared to values from the literature.

(Figure 2–Figure supplement 5E) or loaded (Figure 2–Figure supplement 5F) single molecule data. Thus, we find it unlikely that these stiff mechanics are an artifact of model parameter selection, and instead attribute the high *L_P_* to the experimental conditions and stiff covalent attachments (*Brenner et al., 2016*) required to connect different components in single molecule fluorescence force experiments. Specifically, the incorporation of an SMCC linker connecting the fluorescent molecules to the polypeptide chains likely contributes to the overall mechanics of the chain. The effects of this contribution may be greater for shorter polypeptide sequences, potentially explaining the increased discrepancy between the model and the data in this case. The fit parameters for the different constructs are summarized in Figure 2-Supplementary table 1.

### IX. Comparison to other descriptions of TSMod mechanical sensitivity

In validating the proposed model, we evaluated its ability to recapitulate published fluorescence force spectroscopy measurements from (*Brenner et al., 2016*) of the mechanical response of (GPGGA)_n_ extensible domains flanked by Cy3 and Cy5 fluorescent dyes (Section VIII). Based on their modelling efforts, one of the main conclusions of Brenner *et al.* was that (GPGGA)_n_ behaves as a linear nanospring, not an unstructured polypeptide. This was primarily based on the observation of a linear relationship between average polypeptide end-to-end distance 〈*r_e_*〉 and polypeptide length *N, 〈r_e_〉∼N^1^*. On the contrary, for unstructured polypeptides, the predicted scaling is 〈*r_e_*〉∼*N*^1/2^ (*Dittmore et al., 2011*) or 〈*r_e_〉∼N^3/5^* if excluded volume effects are considered (*Pincus, 1976*). Notably, the calibration model used in this work exhibits 〈*r_e_〉∼N^1/2^* scaling. Two main assumptions differentiated the Brenner calculations from the proposed calibration model. First, the calibration model considers a probability distribution of sensor conformations while Brenner *et al.* performed distance-FRET conversions on an average value. As previously discussed in Section I, premature numerical averaging could lead to numerical artifacts when nonlinear transformations, as are present in distance-FRET conversions, are involved. Secondly, different assumptions were used pertaining to the connectivity of the fluorescent moieties to the polypeptide. Brenner *et al.* used a simple subtraction method which assumes that the fluorescent moiety is located at a constant offset from the extensible domain and *r_e_ = r_c_ — C.* This simple method has previously been reported to lead to numerical artifacts (*Evers et al., 2006a*). In contrast, we used a heuristic *r_e_* to *r_c_* conversion (details in Section III) based on previous simulations of the various physical orientations of TSMod components (*Evers et al., 2006a*).

To determine which of these distinct descriptions is most accurate, we investigated the effects of the assumptions in the two modelling approaches on the predictions of the scaling between 〈*r_e_*〉 and *N.* We first examined what scaling is observed when the proposed calibration model is used to calculate the relationship between 〈*r_e_*〉 and *N* from experimental data. Starting with the published unloaded FRET efficiency measurements of Brenner *et al.* (Figure 2A), the calibration model was used to perform the abovementioned distance-FRET conversions and thus calculate 〈r_e_〉. Plotting this data as a function of *N* yielded a 〈*r_e_〉∼N*^0.57^ scaling (Figure 2–Figure supplement 6A). While we are hesitant to over-interpret this result as it infers power-law relationships from data that varies by less than an order of magnitude, it nonetheless supports the notion that (GPGGA)_n_ mechanics are consistent with unstructured polypeptides. Next, we investigated the prediction of the observed scaling between 〈r_e_〉 and *N* entirely *in silico.* When the calibration model is used to simulate 〈*E*〉 for a collection of TSMods of various lengths *N*, and subsequently the approach of Brenner *et al.* is used to convert these 〈*E*〉 to 〈*r_e_*〉), we then observe an apparent 〈*r_e_*〉∼*N^1^* scaling over the range of polypeptide lengths examined (25 < *N <* 50 residues, Figure 2–Figure supplement 6B). Note that the polymer mechanics model used as the basis of the calibration model predicts 〈*r_e_*〉∼*N*^05^ (Figure 2–Figure supplement 2B). Thus, we conclude that the observation of a spring-like 〈*r_e_*〉∼*N^1^* behavior is a consequence of the assumptions made by Brenner *et al.,* which have previously been shown to lead to poor estimate of 〈*r_e_*〉 (*Evers et al., 2006a; Vogel et al., 2012*). In total, these simulations and data show that the calibration model developed in this work is capable of accurately describing the mechanical properties of unstructured polypeptides, and suggests that extensible domains based on the consensus sequence (GPGGA)_n_ fall within this class.

**Figure 2–Figure supplement 6.**
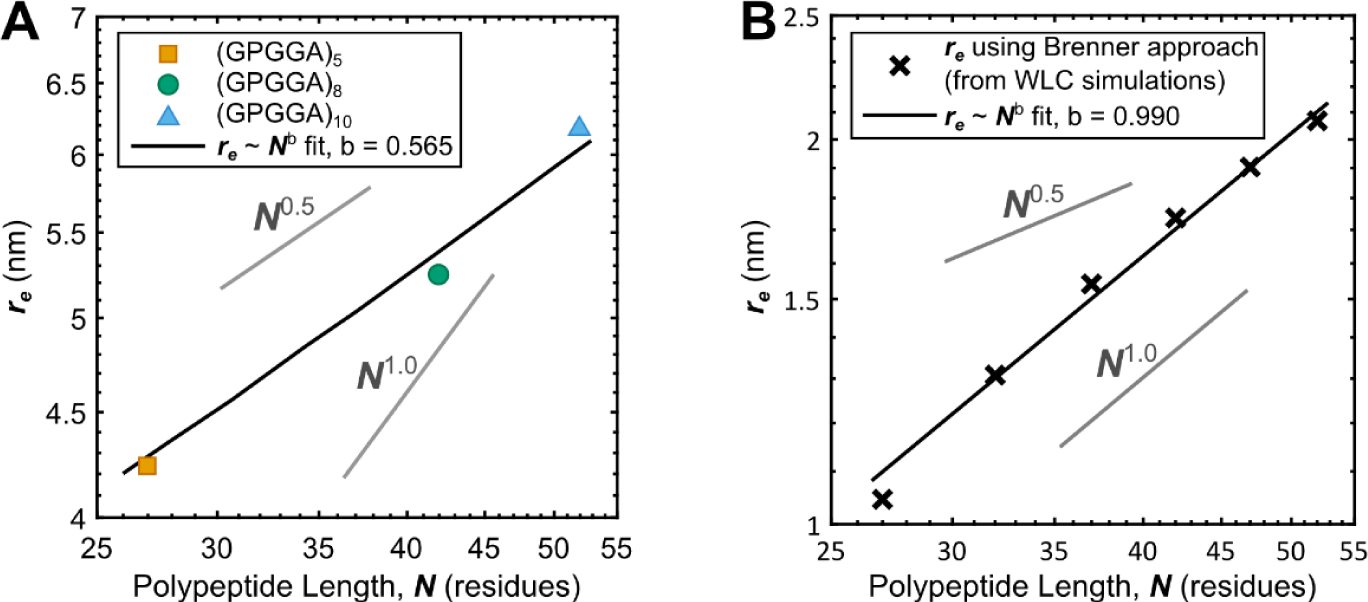
Experimental and theoretical examinations of other models (*Brenner et al., 2016*) of (GPGGA)_n_ mechanical sensitivity. (**A**) Measured relationship between average polypeptide end-to-end distance *r_e_* as a function of the number of residues in the (GPGGA)_n_ extensible domain *N* shows *r_e_∼N^1/2^* behavior characteristic of unstructured polypeptides; data transformed from Figure 2A using the proposed model of TSMod mechanical sensitivity. (**B**) After simulating unstructured polypeptides of various lengths in the context of the WLC model, calculation of *r_e_* using the approach put forth by (*Brenner et al., 2016*) yields an artificial *r_e_∼N^1^*behavior. Note that plots are logarithmic on both axis such that power-law relationships appear as straight lines.

**Figure 2-Supplementary table 1.**
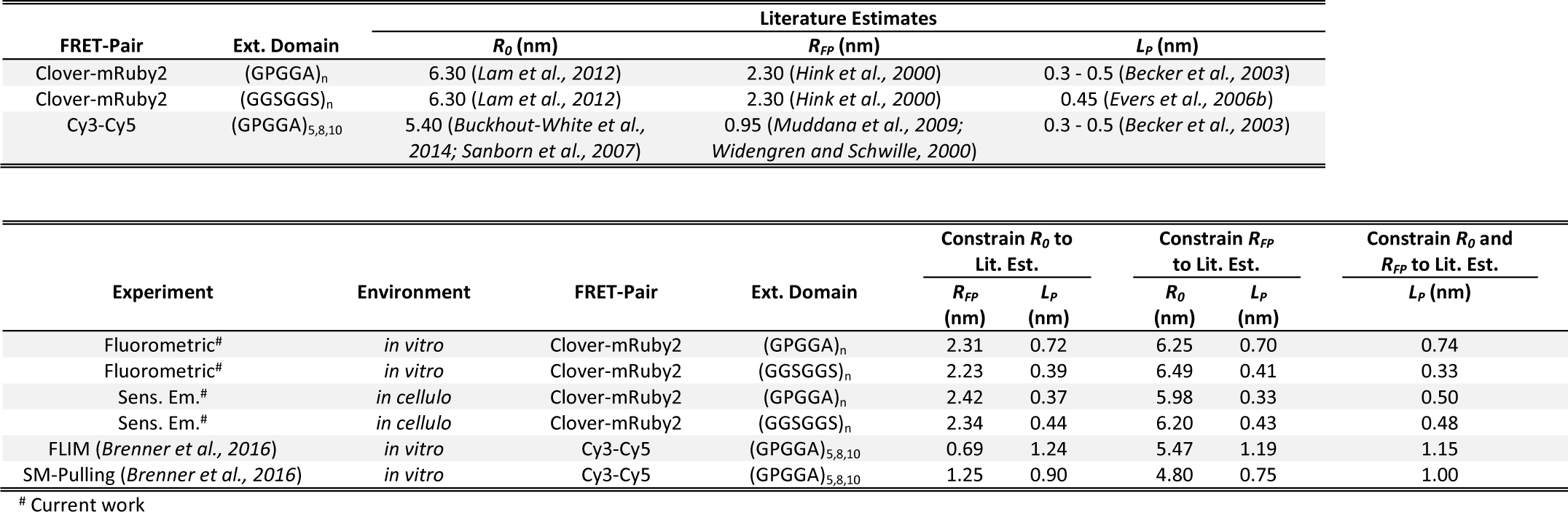
Fit parameters compared to literature estimates of Förster radius (R_0_), fluorescent moiety radius (R_Fp_), and persistence length (L_p_).

| FRET-Pair | Ext. Domain | Literature Estimates |  |  |
| --- | --- | --- | --- | --- |
| | | $R_0$ (nm) | $R_{FP}$ (nm) | $L_P$ (nm) |
| Clover-mRuby2 | (GPGGA)<sub>n</sub> | 6.30 (*Lam et al., 2012*) | 2.30 (*Hink et al., 2000*) | 0.3 - 0.5 (*Becker et al., 2003*) |
| Clover-mRuby2 | (GGSGGS)<sub>n</sub> | 6.30 (*Lam et al., 2012*) | 2.30 (*Hink et al., 2000*) | 0.45 (*Evers et al., 2006b*) |
| Cy3-Cy5 | (GPGGA)<sub>5,8,10</sub> | 5.40 (*Buckhout-White et al., 2014; Sanborn et al., 2007*) | 0.95 (*Muddana et al., 2009; Widengren and Schwille, 2000*) | 0.3 - 0.5 (*Becker et al., 2003*) |

| Experiment | Environment | FRET-Pair | Ext. Domain | Constrain $R_0$ to Lit. Est. | | Constrain $R_{FP}$ to Lit. Est. | | Constrain $R_0$ and $R_{FP}$ to Lit. Est. |
| --- | --- | --- | --- | --- | --- | --- | --- | --- |
| | | | | $R_{FP}$ (nm) | $L_P$ (nm) | $R_0$ (nm) | $L_P$ (nm) | $L_P$ (nm) |
| Fluorometric<sup>#</sup> | *in vitro* | Clover-mRuby2 | (GPGGA)<sub>n</sub> | 2.31 | 0.72 | 6.25 | 0.70 | 0.74 |
| Fluorometric<sup>#</sup> | *in vitro* | Clover-mRuby2 | (GGSGGS)<sub>n</sub> | 2.23 | 0.39 | 6.49 | 0.41 | 0.33 |
| Sens. Em. <sup>#</sup> | *in cellulo* | Clover-mRuby2 | (GPGGA)<sub>n</sub> | 2.42 | 0.37 | 5.98 | 0.33 | 0.50 |
| Sens. Em. <sup>#</sup> | *in cellulo* | Clover-mRuby2 | (GGSGGS)<sub>n</sub> | 2.34 | 0.44 | 6.20 | 0.43 | 0.48 |
| FLIM (*Brenner et al., 2016*) | *in vitro* | Cy3-Cy5 | (GPGGA)<sub>5,8,10</sub> | 0.69 | 1.24 | 5.47 | 1.19 | 1.15 |
| SM-Pulling (*Brenner et al., 2016*) | *in vitro* | Cy3-Cy5 | (GPGGA)<sub>5,8,10</sub> | 1.25 | 0.90 | 4.80 | 0.75 | 1.00 |

<sup>#</sup> Current work

**Figure 2-Supplementary table 2.** Persistence length (*L_p_*) for a variety of published polypeptides. Overall, persistence lengths < 0.2 nm and > 5.0 nm are rarely observed. Synthetic homo-polymers (ex. poly-proline) can achieve larger persistence lengths.

| Polypeptide | $L_P$ (nm) | Ref. |
| --- | --- | --- |
| Titin | 0.2 | (*Kellermayer et al., 1997*) |
| Titin Ig domain | 0.4 | (*Rief et al., 1997*) |
| PEVK domain (in titin) | 0.55 – 0.65 | (*Linke et al., 2002*) |
| PEVK domain (in titin) | 0.92 | (*Sarkar et al., 2005*) |
| Tenascin FN-III domain | 0.56 | (*Oberhauser et al., 1998*) |
| Whole spider silk protein | 0.3 – 0.5 | (*Becker et al., 2003*) |
| Poly-glycine | 0.41 | (*Evers et al., 2006b*) |
| Poly-alanine | 1.76 | (*Evers et al., 2006b*) |
| Poly-proline | 22.0 | (*Evers et al., 2006b*) |
| Poly-proline | 4.5 | (*Schuler et al., 2005*) |
| Panel of short peptides | 0.53 – 1.09 | (*Ohashi et al., 2007*) |
| Elastin | 0.19 | (*Valiaev et al., 2008*) |
| (GGSGGS)<sub>n</sub> | 0.45 | (*Evers et al., 2006b*) |
| (GPGGA)<sub>n</sub> – *in vitro* | 0.74 | This work |
| (GGSGGS)<sub>n</sub> – *in vitro* | 0.33 | This work |
| (GPGGA)<sub>n</sub> – *in cellulo* | 0.5 | This work |
| (GGSGGS)<sub>n</sub> – *in cellulo* | 0.48 | This work |

**Figure 3–Figure supplement 1.**
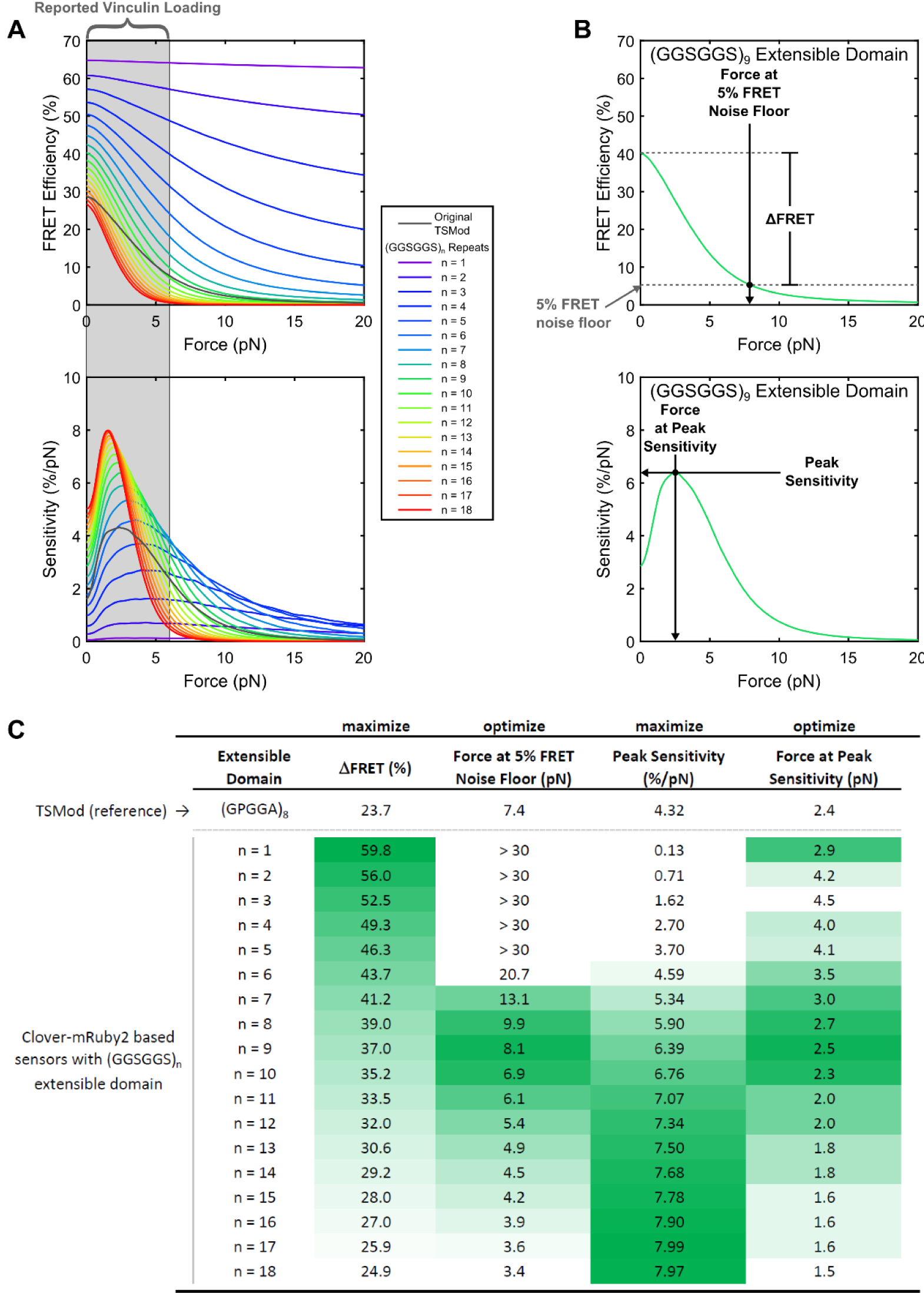
Selection of optimal (GGSGGS)_n_ extensible domain length in a Clover-mRuby2 based TSMod for measuring ∼1-6 pN loads borne by vinculin. (**A**) Predicted force sensitivities of TSMods containing “minimal” Clover-mRuby2 FRET pair and (GGSGGS)_n_ extensible domains (n =1 to 16). (**B**) Schematic of metrics used to quantify and compare different aspects of TSMod mechanical sensitivity, which describe sensor sensitivity (Δ*FRET,* top; *Peak Sensitivity,* bottom) and functional force measurement regimes (*Force at 5% FRET noise floor,* top; *Force at Peak Sensitivity,* bottom) (**C**) Graphical comparison of TSMod mechanical sensitivity properties; green indicates favorable value for a given metric. Nine-repeat (GGSGGS)_n_ extensible domain shows similar force range as original TSMod (column 2) as well as a good balance between Δ*FRET* (column 1) and *Peak Sensitivity* (column 3), which are inversely proportional.

**Figure 3–Figure supplement 2.**
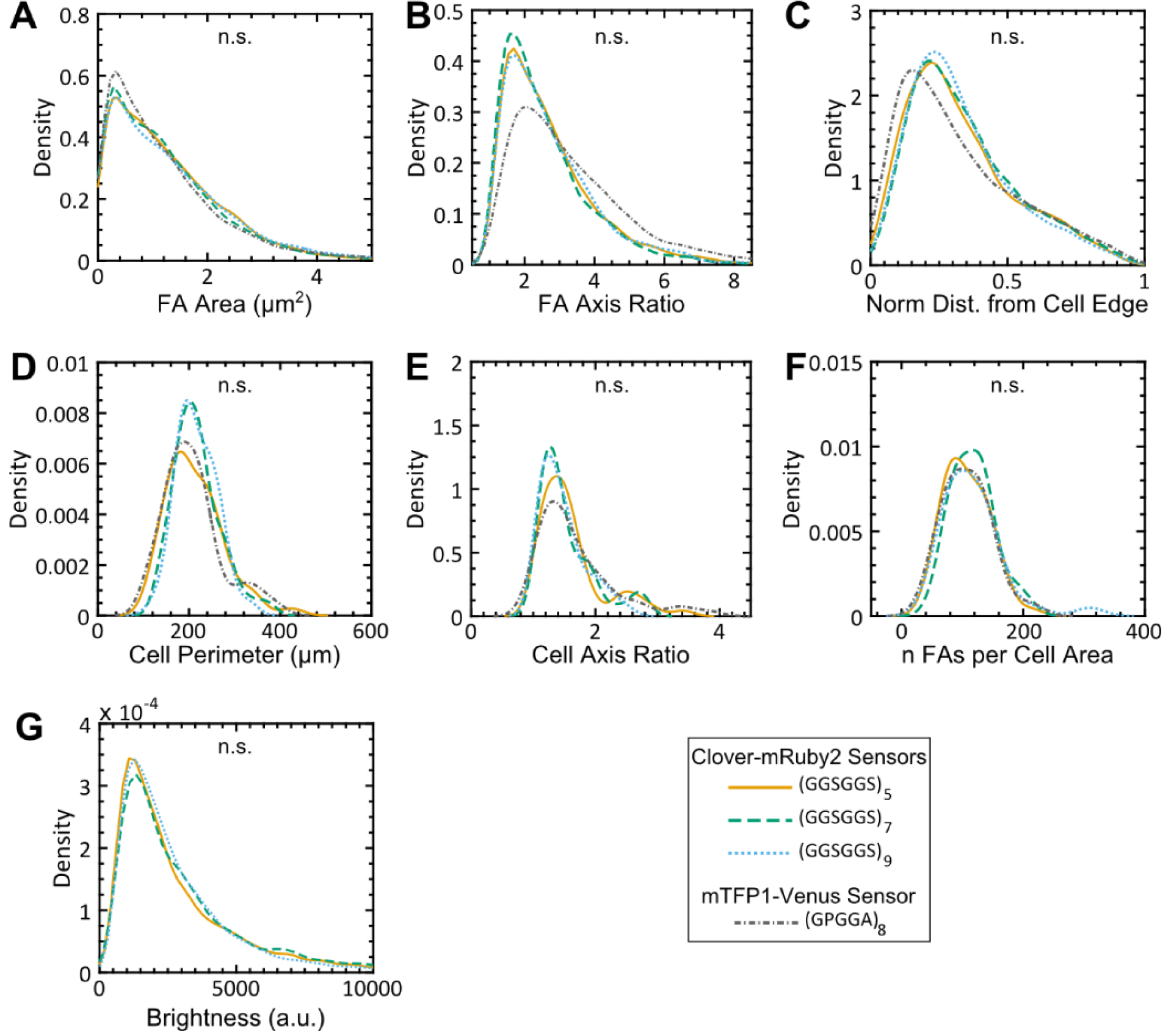
FA morphologies, cell morphologies, and sensor localization to FAs are indistinguishable between different versions of VinTS. (**A-G**) Vin-/- MEFs expressing various version of VinTS show indistinguishable FA area (**A**), FA axis ratio (**B**), subcellular distributions quantified as normalized distance from cell edge (**C**), cell perimeter (**D**), cell axis ratio (**E**), number of FAs normalized by cell area (**F**), sensor localization to FAs in terms of mean acceptor intensity (brightness) (**G**). Different versions of the tension sensor include those constructed with minimal Clover-mRuby2 modules containing three distinct extensible domains, namely (GGSGGS)_5_ (n = 54 cells), (GGSGGS)_7_ (n = 40 cells), (GGSGGS)_9_ (n = 33 cells), along with the original TSMod containing a (GPGGA)_8_ extensible domain (n = 48 cells), each from 3 independent experiments; n.s. not significant (p ≥ 0.05), ANOVA. See **Supplementary file 3** for exact p-values.

## Figure supplements for Figure 4

### Figure 4-Supplementary note 1: Examining extension-based control of vinculin loading with a structural model of a FA

Experiments that showed extension-based control of vinculin loading (Figure 4) leveraged three tension sensors with unique force-extension relationships, which were dictated by the unique mechanical responses of the extensible (GGSGGS)_n_ polypeptide domains. Seeking to gain a better understanding of what kinds of physical interactions might give rise to force-controlled versus extension-controlled loading of different FA components, we developed a simple structural model of a single FA. The FA structural model consists of 170 elements each of which is described as a Hookean spring and can be conceptualized as either a “sensor” element or an alternative “linker” element. Based on the reported stratified organization of layers of proteins within FAs (*Kanchanawong et al., 2010*), we arranged the two elements in two parallel layers (Figure 4–Figure Supplement 2A). However, even with this simplified geometry, numerous scenarios are possible. Key variables include the relative stiffness and relative abundance of each element, as well as whether a bulk force input or a bulk extension input is provided to the structure. Therefore, the focus of these modeling efforts was to determine how the force-controlled or extension-controlled loading of individual FA proteins might be impacted by each of these variables in a variety of scenarios.

To examine a single possible physical scenario, structural models were considered in groups of three (Figure 4–Figure Supplement 2B), since experimental evaluation of force-based versus extension-based control similarly involved three sensors with distinct mechanical properties (Figure 4). Within each grouping of three, the stiffness of the linker elements, *k_L_*, is held constant and only the stiffness of the sensor element was varied (*k*_*S*1_, *k*_*S*2_, or *k*_*S*3_). The stiffness values used for *k*_*S*1_, *k_*S*2_,* and *k_*S*3_* were selected to approximate the relative differences in stiffness between the (GGSGGS)5,7,9 extensible domains used in experiments and were estimated from linear fits to their predicted force-extension curves (Figure 4–Figure Supplement 1). Figure 4–Figure Supplement 2B depicts a single scenario in which equal numbers of sensor and linker elements (*N_s_* = *N_L_*) with comparable average stiffness (〈*k_Sj_*〉 = *k_L_*) are loaded by a bulk extension input (*δ*_0_) as an illustrative example.

Within any particular geometry, we evaluated the extent of force-controlled versus extension-controlled loading of the three distinct sensor elements. Following bulk loading of the structure, which in the illustrative case corresponds to a constant extension *δ*_0_,we solve for the extension of and force across the sensor elements in each of the three assemblies (*δ*_*sj*_ and *F*_sj_, respectively). The extent of extension-controlled versus force-controlled mechanical behaviors is then evaluated by comparing the coefficient of variations, which is defined as standard deviation divided by the mean and abbreviated *CV* here, of the forces *CV(F_sj_*) and extensions *CV(S_sj_*) experienced by these three different stiffness sensor elements. A control metric quantitatively relating the two magnitudes of variation is defined as the log2 ratio of the two coefficients of variation:

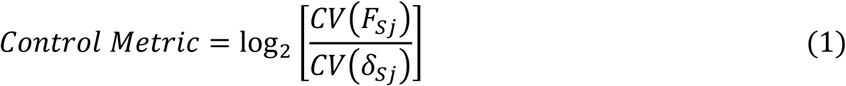

**Figure 4–Figure supplement 1.**
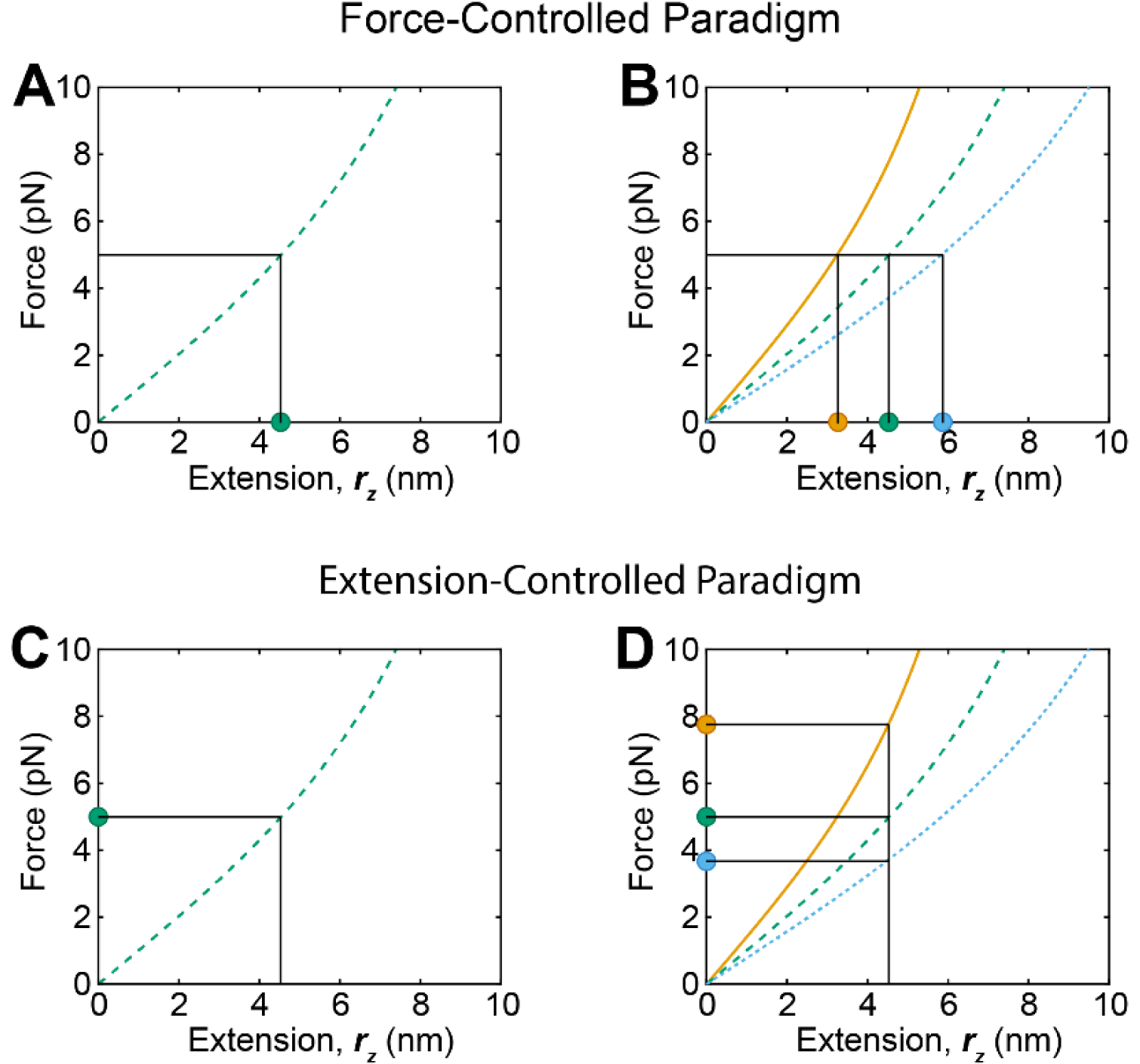
Schematic depiction of force-extension relationships and potential force- and extension-control paradigms. (**A, B**) A force controlled paradigm cannot be detected if only a single sensor is used (**A**), but can be discerned if multiple sensors report the same forces, but distinct extensions (**B**). (**C, D**) An extension controlled paradigms also cannot be detected if a single sensor is utilized (**C**), but can be uncovered if multiple sensors report the same extension and therefore distinct forces across a protein of interest (**D**). Curves are predictions of the force-extension curve of the WLC model with persistence lengths of 0.5 nm and contour lengths corresponding to the (GGSGGS)5,7,9 linkers utilized in this study (orange, green, and blue, respectively).

In an extension-controlled assembly the forces will vary more than the extensions *CV(F_sj_) > CV(δ_sj_*), resulting in a large and positive control metric, while the opposite will be true for a force-controlled assembly where *CV(F_sj_) < CV(δ_s_j*). In the balanced scenario depicted in Figure 4–Figure Supplement 2B, which contains equal numbers of sensor and linker elements of identical average stiffness, we observe neither extension nor force control following a bulk extension input to the assemblies. Thus, the control metric is close to zero.

In this framework, we investigated the effects of both bulk force (*F_0_*) and extension (*δ*_0_) inputs to a variety of stratified structures comprised of variable abundances and stiffnesses of the two types of springs (Figure 4–Figure Supplement 2C). To accomplish this, the above calculations were repeated while the relative number of springs and *k_L_* were varied. As we are most interested in relative rather than absolute numbers of elements and element stiffness, the relative numbers and mechanics of the sensor and linker elements are defined as:

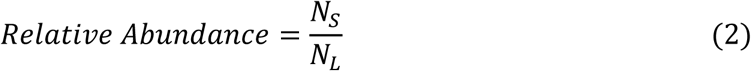

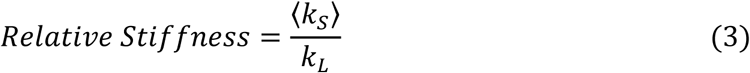

The number of sensor and linker elements (*N_s_* and *N_L_*, respectively) and the stiffness of the linker element (*k_L_*) were set such that the relative abundance and relative stiffness of the two elements varied between 2^−4^ and 2^4^ Note that when calculating the relative stiffness, the three distinct spring constants for the three sensor elements were averaged together. This definition is appropriate as the variation of the stiffness of the sensors springs (*k_s_j*) is significantly smaller (∼30% variation) than the range of *k_L_* that was evaluated. These normalized parameters also allow us to draw conclusions that are independent of the absolute values of *k* and *N* that are simulated.

With a bulk force input (*F*_0_), simple relationships that obey Hooke’s Law are observed, and *F_0_* is evenly distributed across each sensor element:

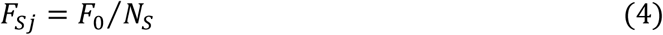

where *F_Sj_* refers to force observed in one of three group models. Subsequently, the force *F_sj_* dictates the extension *δsj* of individual sensor elements following:

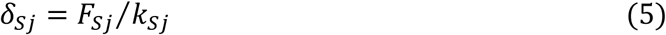

In this mechanical assembly, force *F_sj_* remains constant regardless of the stiffness of the sensor element *k_Sj_*, while extension *δ_sj_* of the sensor element scales inversely with its stiffness *k_sj_*. This simple solution, where forces are constant and extensions change with sensor element stiffness, is a prime example of an assembly exhibiting force-based control.

In response to a bulk extension input (*δ*_0_), more complex behaviors are observed. We must first determine the total force *F* across the assembly following Hooke’s Law for springs in series:

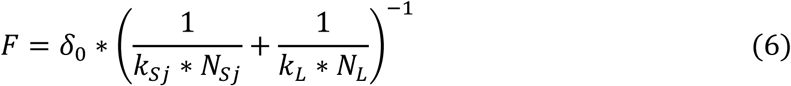

where the effective spring constant for the whole assembly is given by:

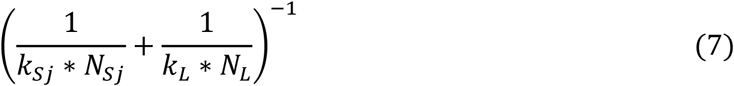

Calculating *F_Sj_* and *δ_sj_* as before (Eq. 4 and 5), it becomes apparent that both *F_S_j* and *sj* will change depending on the relative abundance and stiffness of the sensor and linker elements.

By performing this operation over a variety of relative abundance and stiffness of the sensor and linker elements, we begin to understand what kinds of structural arrangements could give rise to force-controlled versus extension-controlled loading of FA components (Figure 4–Figure Supplement 2D). Specifically, the assemblies most biased toward extension-controlled behaviors involve linker elements that are much stiffer and/or in greater molecular abundance as compared to sensor elements. However, in the opposite scenario, force-controlled loading could be observed even following a bulk extension input. The sensitivity of the control paradigm to sensor and linker element stiffness and abundance indicates that force-controlled and extension-controlled scenarios are likely mutable and highly dependent upon protein, cellular, and mechanical contexts. In our experimental context, the measured variance in forces and extensions reported by the three sensors (Figure 4) indicated a control metric ∼1.9 (Figure 4–Figure Supplement 2D, asterisk). These measurements indicate, at least for MEFs adhered to fibronectin-coated glass substrates, vinculin exhibits predominantly extension-controlled behavior.

The model also predicts the relationship between sensor element stiffness *k_sj_* and the force borne by the sensor element *F_sj_*. Importantly, for various values of relative abundance or stiffness of the sensor and linker elements, the various control regimes correspond to distinct relationships between sensor element stiffness *k_sj_* and force *F_sj_*. To illustrate this, we investigated three scenarios: a region in the extension-controlled regime consistent with our observations of vinculin, a region where neither force- nor extension-based control is favored, and a region in the force-controlled regime (Figure 4–Figure Supplement 2D, dashed contour lines). The extension-controlled regime is associated with a linear relationship between sensor element stiffness and force, while the force-controlled regime predicts a constant relationship between these variables, and the intermediate regime corresponds to a relationship between these two limits (Figure 4–Figure Supplement 2E). The magnitudes of the forces depicted on the y-axis must be considered qualitatively, as they are affected by the magnitude of the bulk extension input (*δ*_0_). Regardless, it is clear that “minimal” Clover-mRuby2 based VinTS with 5, 7, and 9 repeats of a (GGSGGS)_n_ extensible domain (*control metric* = 1.9) clearly exhibit a linear relationship between the forces experienced by each sensor and the estimated spring constants, consistent with being in an extension-controlled regime (Figure 4–Figure Supplement 2E, filled points). Thus, this simple structural model accurately reproduces two distinct features of the mechanical responses observed in the suite of vinculin tension sensors.

**Figure 4–Figure supplement 2.**
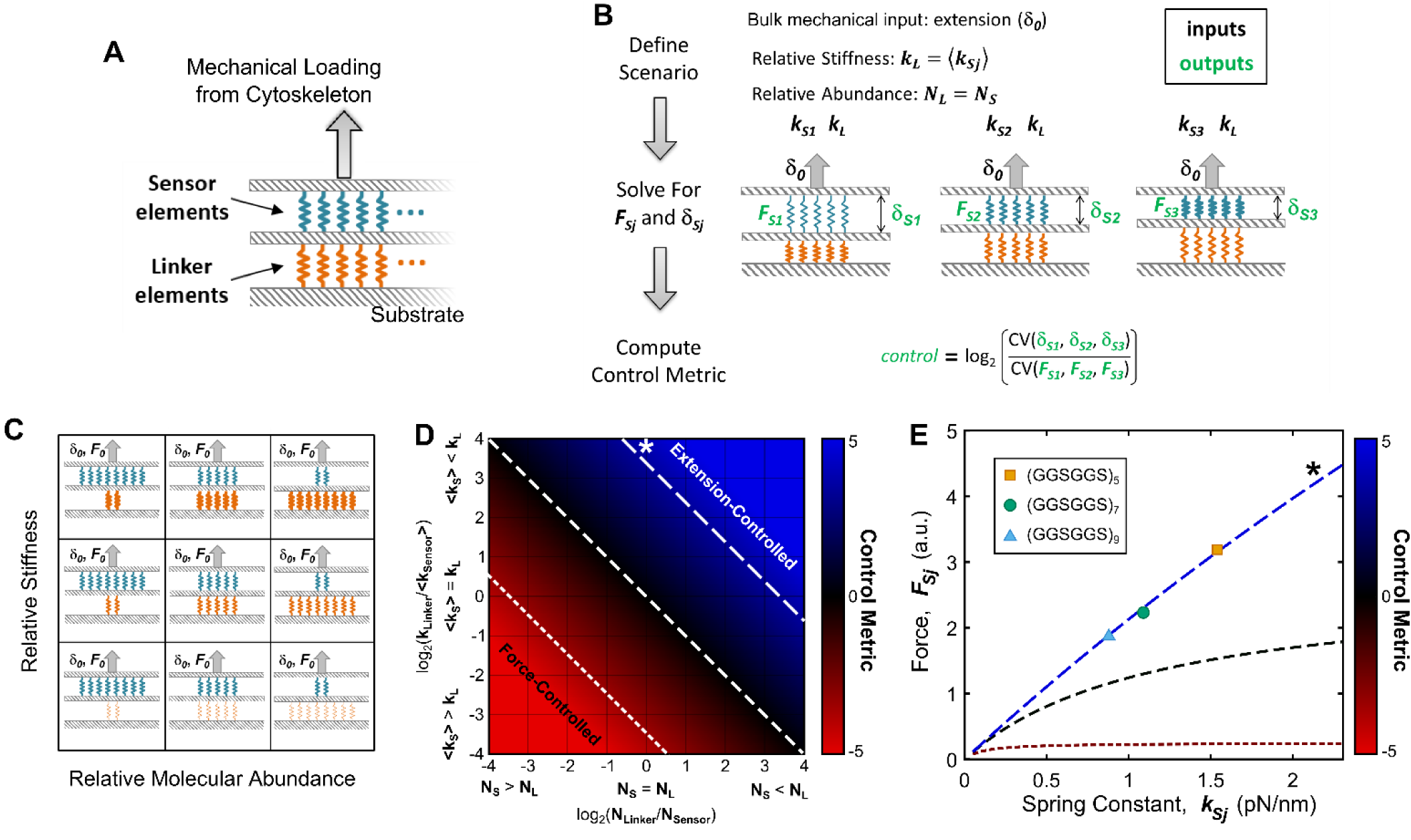
Various structural arrangements within FAs could lead to force-based or extension-based control. (**A**) FA structural model consists of two parallel layers of proteins, which can be conceptualized as either “sensor” (blue) or “linker” (orange) elements. (**B**) Mimicking what was done in experiments (Figure 4), simulations involved three sensor elements with distinct stiffnesses (*k_s1_, k_S2_, k_S3_*), arranged in a stratified fashion with a single linker element (stiffness *k_L_*), loaded by a bulk extension (or force) input. The forces and the extensions experienced by each sensor element were calculated, and a control metric relating the relative variation in forces to the variation in extensions was calculated (see **Figure 4-Supplementary note 1** for details). (**C**) Schematic depiction of parameter space examined using this simple structural model of FAs, wherein the relative number of the sensor and linker element is varied (x-axis) along with their relative stiffness (y-axis); thicker springs indicate stiffer mechanics. (**D**) Summary of results from simulations quantifying force-controlled versus extension-controlled loading of the sensor element. The Control Metric describes the ratio of variation in forces to the variation in extensions experienced by the sensor elements and will be positive for force-controlled situations and negative for extension-controlled situations. Regions of relative stiffness and abundance are highlighted where, following a bulk extension input, force-controlled loading of the sensor element (sensor element is stiff and in relatively high abundance) or extension-controlled loading of the sensor element (sensor element is soft and/or in relatively low abundance) is observed. (**E**) FA structural model predictions of the relationship between sensor element stiffness (spring constant) and force, which are linear for the extension-controlled regime (asterisk), and increasingly flat for balanced and force-controlled regimes; dashed contour lines in panel (**D**) correspond to dashed force-stiffness relationships in panel (**E**).

Together, these model predictions indicate that the extension-controlled loading of individual FA components is most likely to be observed in response to a bulk extension input, will be observed for soft elements in relatively low abundance, and will result in a linear relationship between sensor element stiffness and load bearing capacity, as has been observed experimentally for vinculin. Determining whether other load-bearing proteins, as well as vinculin in other cellular contexts (ex. three-dimensional matrices), are subject to extension-control will be critically important. By developing a rational-design approach for the creation of calibrated FRET-based tension sensors, we uniquely enable and expedite these types of investigations.

## Figure supplements for Figure 5

**Figure 5–Figure supplement 1.**
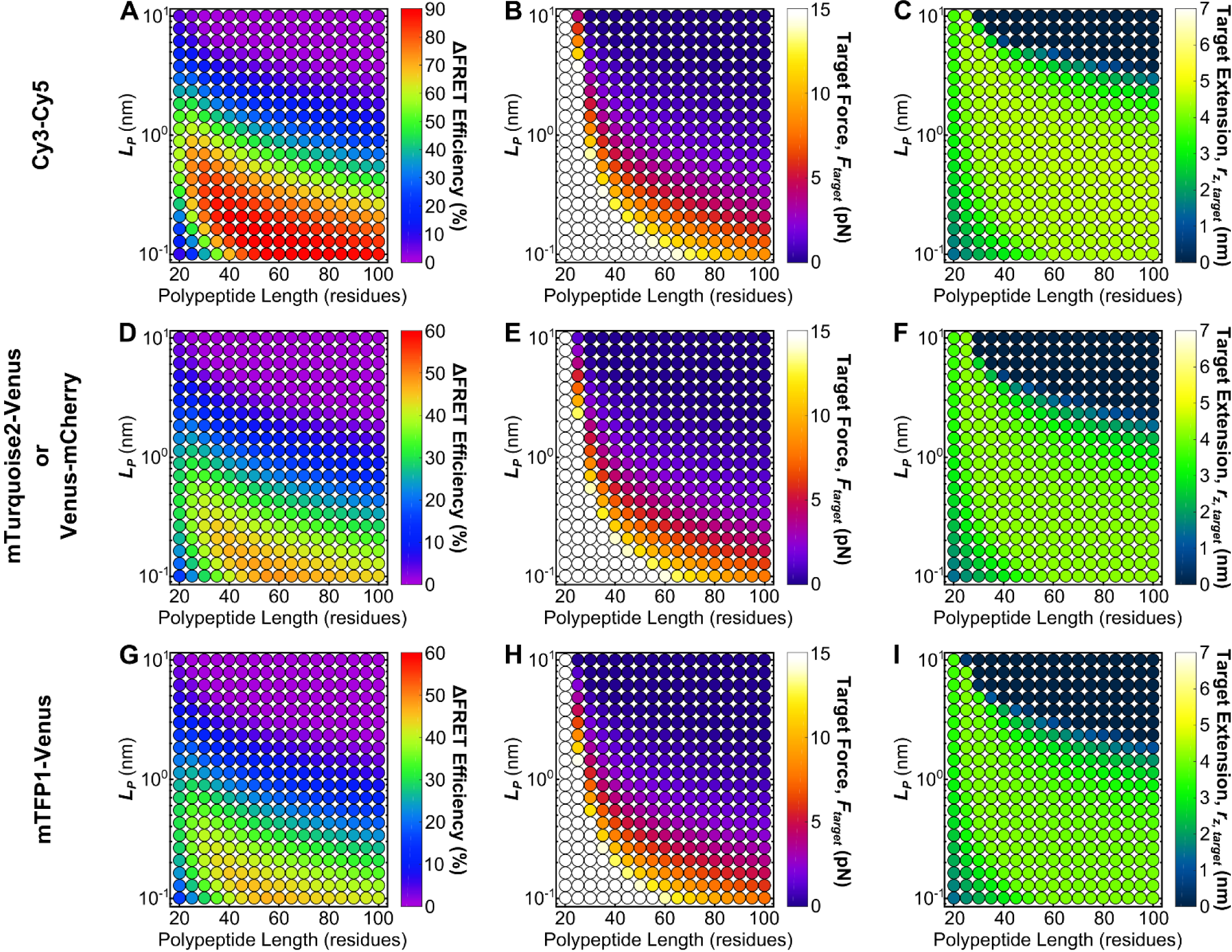
Roadmaps to guide the rational design of FRET-based molecular tension sensors for some commonly used FRET-pairs. Parameter space highlighting the predicted Δ*FRET* at the 5% FRET noise floor (**A, D, G**), as well as corresponding target force, *F_target_* (**B, E, H**), and target extensions *r_z,target_* (**C, F, I**), for a variety of Cy3-Cy5 (**A-C**), Venus-mCherry or mTurquoise2-Venus (which have identical *R_0_*) (**D-F**), and mTFP1-Venus (**G-I**), based sensors with a variety of polypeptide lengths (x-axis), and mechanical properties (y-axis). Each point represents a single potential tension sensor design.

## Supplemental files

**Supplementary file 1.**
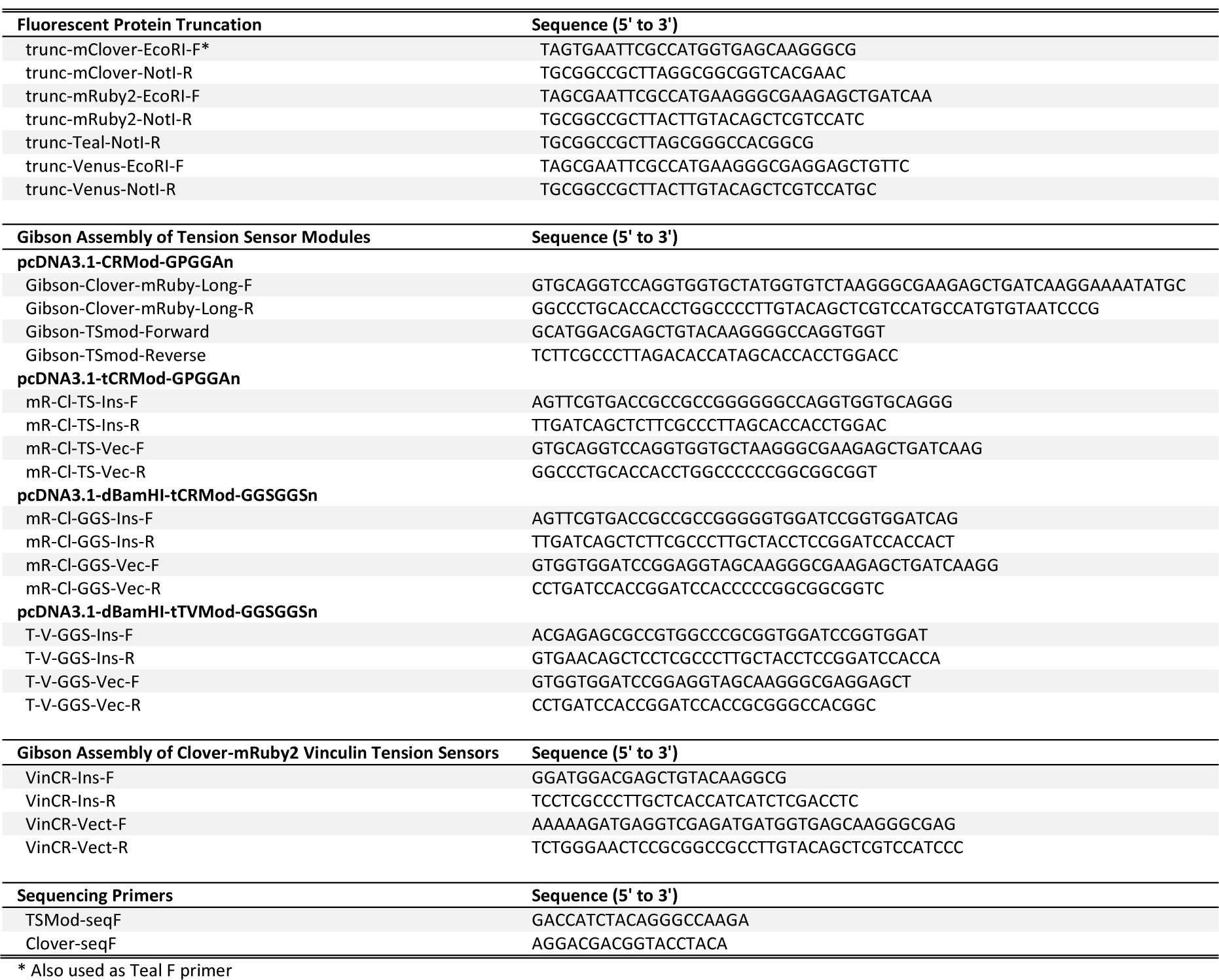
Primers used in this study.

**Supplementary file 2.**
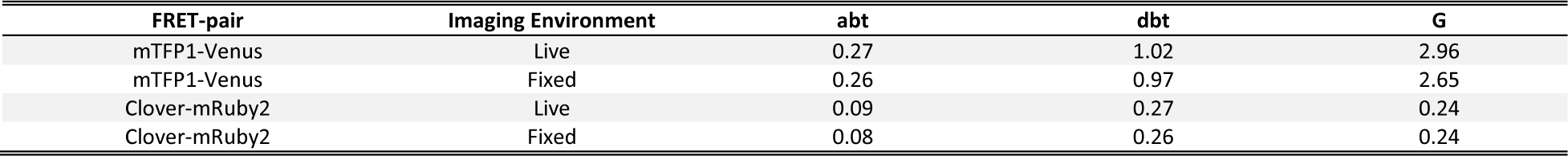
Spectral bleed-through coefficients and G factors for mTFP1-Venus and Clover-mRuby2 based tension sensor FRET efficiency calculations.

**Supplementary file 3.**
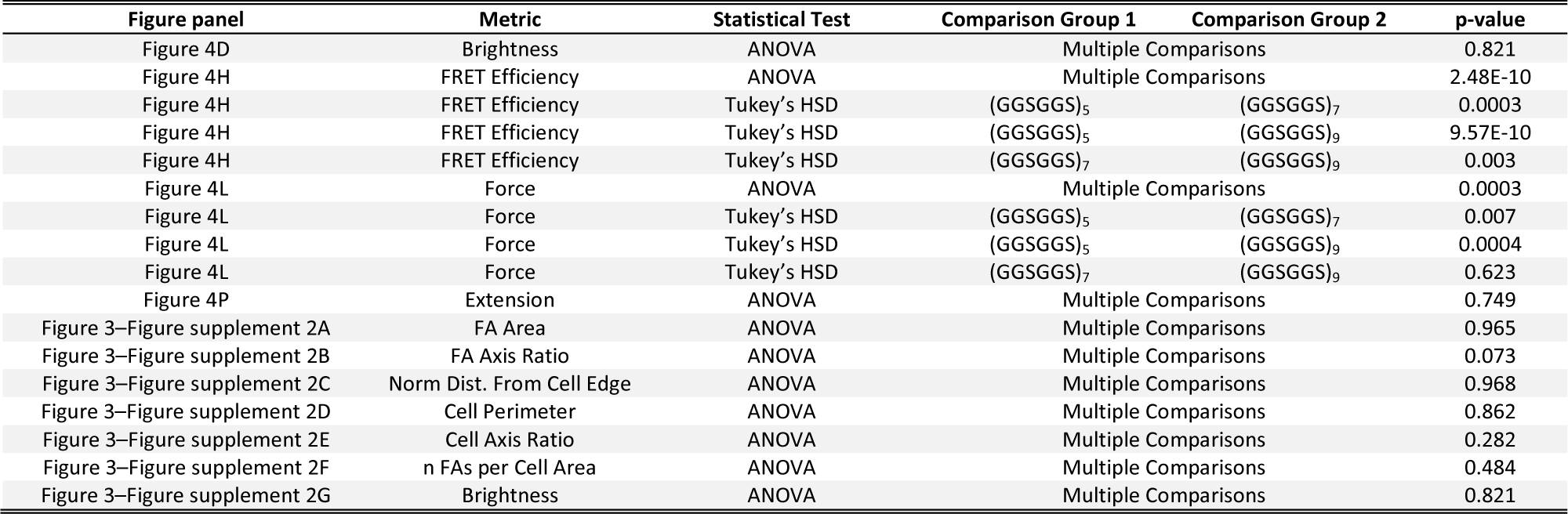
Statistical test details including exact p-values for ANOVAs and post-hoc tests.

